# Comparative effects of proton and photon irradiation on the molecular and cellular profiles of triple-negative breast cancer: the crucial impact of VEGFC on tumor microenvironment remodeling

**DOI:** 10.1101/2024.08.19.608614

**Authors:** Saharnaz Sarlak, Delphine Marotte, Florent Morfoisse, Alessandra Pierantoni, Jessy Sirera, Meng-Chen Tsai, Marie Vidal, Joël Hérault, Barbara Garmy-Susini, Jérôme Doyen, Frédéric Luciano, Gilles Pagès

**Affiliations:** University Cote d’Azur (UCA), Institute for research on cancer and aging of Nice (IRCAN), CNRS UMR 7284; INSERM U1081, Centre Antoine Lacassagne, France; I2MC, Université de Toulouse, Inserm UMR 1297, UPS, 31000 Toulouse, France; Institut Méditerranéen de Protonthérapie—Centre Antoine Lacassagne, Fédération Claude Lalanne, Nice, France

**Keywords:** TNBC, VEGFC, X and P irradiation, resistance, anti-VEGFC antibodies

## Abstract

Metastatic triple-negative breast cancers (TNBC) are among the most aggressive types of breast cancer and are often treated with adjuvant radiotherapy and chemotherapy. Despite initial efficacy, relapses are common, leading to poor prognosis. Understanding the response of tumor microenvironment to radiotherapy is crucial, particularly comparing photon (X) and proton (P) radiotherapy due to proton radiation’s reduced side effects.

**Methods:** We investigated the effects of single and multiple X and P irradiations on various cell types within the tumor microenvironment, including vascular and lymphatic endothelial cells, fibroblasts, and TNBC tumor cells. VEGFC, a key factor in lymphatic vessel formation and metastasis, was a primary focus. We used protein arrays to evaluate the effects of irradiation and examined the impact of VEGFC inactivation on the sensitivity to X and P radiation. Additionally, we tested tumor-forming capabilities of irradiated cells and assessed the impact of genetic or therapeutic VEGFC inhibition on TNBC growth. Transcriptomic and proteomic analyses further characterized the differences between X and P tumors, providing deeper insights into their distinct molecular profiles.

**Results:** Both X and P irradiations caused a transient increase in VEGFC levels, along with other pro-angiogenic, pro-lymphangiogenic, and pro-fibrotic factors, such as angiopoietin 2, artemin, endostatin, IGFBP2, serpinE1, PDGFA, and DPPIV. Endothelial cells exposed to multiple rounds of radiation showed enhanced proliferation but lost the ability to form pseudo vessels, indicating an endothelial-mesenchymal transition. Tumor cells lacking VEGFC were more sensitive to radiation, and anti-VEGFC antibodies significantly suppressed TNBC cells’ proliferation, both naïve and multi-irradiated. Tumor xenografts formed by multi-irradiated cells grew larger in nude mice, particularly following proton irradiation, while X-irradiated tumors exhibited a more pro-lymphangiogenic phenotype compared to P-irradiated tumors.

**Conclusions:** Our findings show that while P multi-irradiated TNBC cells form larger tumors, X multi-irradiated tumors are more aggressive, with elevated expression of genes linked to angiogenesis, lymphangiogenesis, and endothelial-mesenchymal transition. Targeting VEGFC during photon or proton radiotherapy could reduce metastasis and improve TNBC prognosis.

## Introduction

The treatment landscape for breast cancer (BC) has seen substantial advancements over the past decades; however, a specific subset, known as triple-negative breast cancer (TNBC), continues to pose significant clinical challenges [1]. The nomenclature “triple-negative” arises from the absence of reliance on estrogen and progesterone hormone receptors, as well as the HER2 receptor for their growth. TNBCs are characterized by a notably aggressive growth rate and heightened metastatic potential [2].

BC, in general, are distinguished by robust vascularization, alongside the presence of an alternative network of lymphatic vessels that facilitates accelerated metastatic dissemination. Consequently, the identification of the sentinel lymph node plays a crucial role in the diagnostic arsenal, aiding in the classification of tumors as localized or metastatic at different stages.

The standard therapeutic approach for BC adopts a comprehensive, multi-modal strategy that typically begins with surgery and integrates adjuvant radiotherapy and chemotherapy. This holistic treatment regimen is designed to address both local and systemic aspects of the disease, contributing to improved outcomes for individuals with BC.

For hormone-sensitive and HER2-dependent tumors, targeted therapies play a pivotal role. This includes anti-aromatase or anti-estrogen compounds for hormone-sensitive BC [3, 4], and specific antibodies targeting HER2 such as trastuzumab and pertuzumab, along with antibody-drug conjugates like trastuzumab emtansine or trastuzumab deruxtecan for HER2-dependant BC [5].

However, for TNBC, aggressive and traditional chemotherapies, such as taxanes or doxorubicin, remain the standard of care, often accompanied by significant toxic side effects. Recently, the utilization of small molecule inhibitors [6] as well as immunotherapy have emerged as a promising strategy for TNBC treatment, with the combination of atezolizumab (Anti-PDL1) [7, 8] or pembrolizumab (anti-PD1) [9, 10] and Nab-paclitaxel showing efficacy. The utilization of the anti-PD1 antibody pembrolizumab in a neoadjuvant setting has also demonstrated significant promise [11, 12]. Neoadjuvant therapy involves administering treatment before surgery, with the aim of reducing the size of tumors or eliminating micro metastases. Pembrolizumab, by targeting the PD1 pathway, contributes to enhancing the ability of immune system to recognize and attack cancer cells. Neoadjuvant pembrolizumab may not only facilitate the management of existing tumors but also holds the potential to enhance the overall effectiveness of subsequent therapeutic interventions.

Advancements in radiotherapy have been significant over the last few decades, with more specific and precise irradiation procedures, notably using advanced technologies like the CyberKnife apparatus [13, 14]. Furthermore, the introduction of proton-based therapy, as opposed to photon-based radiotherapy, holds great promise due to its efficacy and more precise irradiation, which helps narrow the radiation field and limit side effects. Nevertheless, the financial implications linked to proton therapy remain a substantial concern. To address this, ongoing clinical trials are actively underway to substantiate and demonstrate the comparative advantages of proton (P) therapy in contrast to traditional photon (X) irradiation, as highlighted by recent research [15]. These trials aim to provide valuable insights into the cost-effectiveness and clinical efficacy of proton therapy, contributing to informed decision-making within the medical community. Despite the effectiveness observed in various treatment schedules, recurrence remains a significant challenge, particularly in the case of the highly aggressive TNBC. Notably, across diverse treatment regimens, irradiation schedules emerge as a common factor in BC management.

Building on our previous findings regarding head and neck tumors [16], where both P and X irradiation induced stress on tumor cells, ultimately leading to cell death, we extend this perspective to the BC context. Contrary to conventional beliefs, irradiation is not only a stressor for targeted tumor cells but also impacts normal cells within the tumor microenvironment and at the tumor margin. Viewing irradiation as a stressor capable of modifying the genetic program of both normal and tumor cells, we hypothesize that, under certain conditions, irradiation may exhibit a protumor effect. This effect may involve the shaping of tumor cells that have not received the maximal dose of P or X, but also altering the genetic landscape of normal cells to release growth factors or inflammatory cytokines beneficial for a protumor environment. Building on our previous work, where P demonstrated equivalent therapeutic efficacy on tumor cells but distinctively shaped the production of the pro-lymphangiogenic factor VEGFC [16], we conducted a comprehensive analysis comparing the relative effects of P and X on various components of the tumor microenvironment. This included tumor cells, vascular and lymphatic endothelial cells, and fibroblasts. Our findings reveal that P multi-irradiated TNBC cells (used to simulate radioresistant cells) form larger tumors compared to X multi-irradiated cells. However, at the molecular level, X multi-irradiated tumors display greater aggressiveness, with higher expression of genes related to angiogenesis, lymphangiogenesis, and endothelial-mesenchymal transition (Endo-MT). In this study, the pro-lymphangiogenic factor VEGFC emerges as a key player in this protumor effect. These results underscore the complexity of irradiation impacts on the tumor microenvironment and highlight the need for a nuanced approach in tailoring treatment strategies for TNBC subtypes.

## Methods

### Cell Lines and Culturing Conditions

Human triple negative breast cancer cell lines (MDAMB231, BT549, CAL51, E0, 4T1) and telomerase-immortalized microvascular endothelial (TIME) were purchased from the ATCC. These cells were cultured in DMEM (1X) + GlutaMAX 4.5 g/L D-Glucose (Cat. N. 10566016, Gibco®) supplemented with 10% Fetal Bovine Serum (FBS) (Cat. N. 10270106, Gibco®), 1% MEM Non-Essential Amino Acids Solution, (Cat. N. 11140050m, Gibco®) and 1% penicillin-streptomycin (Gibco®). Human Dermal Lymphatic Endothelial Cells (HDLEC) (Cat. N. C-12216) and Human Umbilical Vein Endothelial Cells (HUVEC) (Cat. N. C-12203) pooled were purchased from the Promocell (Heidelberg, Germany). LEC cells cultured in Endothelial Cell Basal Medium w/o Glutamine from PELObiotech (Cat. N. PB-BH-100-9806). TIME and HUVEC cultured in Vascular Cell Basal Medium from ATCC (Cat. N. PCS-100-030™). All the cells maintained at 37 °C in humidified atmosphere at 5% CO2.

*VEGFC^KO^ clones*: The *VEGFC* gene was knocked-out in wild type MDAMB231 and BT549 cells by the CRISPR-Cas9 technique [17]. Briefly, a human *VEGFC* target oligonucleotide (5′-GAGTCATGAGTTCATCTACAC-3′) was cloned into the pX330-U6-Chimeric_BB-CBh-hSpCas9 vector (Addgene plasmid # 42230). Two *VEGFC*^KO^ clones were obtained by PEI transfection (Tebu Bio, Le-Perray-en-Yvelines, FRANCE) of the resulting vector into MDAMB231 or BY549 cells and further selection on 5 µg/ml puromycin (InvivoGen, Toulouse, France), for 10–15 days. Control cells (Ctl) were obtained by transfection of WT-MDAMB231 or WT-BT549 cells by an empty pX330 vector and puromycin selection. The mutations leading to *VEGFC* invalidation were revealed, for each clone, by genomic DNA sequencing, using the following primers: Sense, 5′-TTGTGTTAGGGAACGGAGCAT-3′; Antisense, 5′-AGAACCAGGCTGGCAACTTC-3′ **(Supp. Table S1)**. Effectiveness of *VEGFC* invalidation was confirmed by ELISA assay of VEGFC production.

### Cell irradiations

Hundred fifty thousand cells were seeded in the 12 cm^2^ tissue culture flasks, 24 h prior to the irradiations. Proton irradiation was done at Centre Antoine Lacassagne, Nice (CAL) using 63 MeV Cyclotron MEDICYC and Photon irradiation was done using a Faxitron cabinet X-ray irradiator (160kV-6.3 mA; Edimex, Le Plessis-Grammoire, FRANCE) in 2, 4 and 8 Gy.

### Cell proliferation

Cells were seeded in six-well plates in triplicates and were counted every day or every 48 h for 6 to 8 days, using a Coulter counter (Villepinte, FRANCE).

### Colony formation assay

Cells (500 cells per condition) were resuspended with their own corresponding medium and seeded in triplicate in 6-well plates. Colonies were detected two weeks after seeding. They were fixed at room temperature for 20 min with 3% paraformaldehyde (PFA; Electron Microscopy Sciences) and stained with Crystal Violet. The number and size of the clones were analyzed using ImageJ software.

### Transmigration

The transmigration assay based on chemotaxis was performed by inserting a transwell polyester membrane filter with 8μm pores polycarbonate membrane (Falcon® Cell Culture Inserts. Product number: 353097) in 24-well culture plates. Total of 150 000 serum starved MDAMB231, BT549 and TIME cells were resuspended in 300μL of the serum free corresponding media seeded in upper chamber of Transwell. To chemoattract the cells, corresponding medium containing 10% FBS was placed in the lower chamber. After 24h of incubation, the migrated cells on the bottom side of the membrane were counted. Each assay was performed in triplicate wells. Cells were counted using the ImageJ software (NIH; Bethesda, MD, USA).

### Enzyme-linked immunosorbent assay (ELISA)

The conditioned media of cells 48 h post seeding were collected, and the corresponding cells were counted for data normalization. VEGFC was quantified from supernatants with the R&D Systems Human VEGFC Duoset ELISA kit (Minneapolis, MN, USA), according to the manufacturer’s recommendations.

### RNA extraction and cDNA synthesis

Total RNA was isolated from cells by using RNeasy Mini Kit (Quiagen) according to the manufacturer’s instructions. Total RNA was quantified spectrophotometrically, and 1 µg was treated with 1U of DNAse RNAse-free (Quiagen). cDNA synthesis was performed by using the QuantiTect Reverse Transcription Kit (QIAGEN), according to the manufacturer’s instructions. For oligo sequences, see Supplementary Materials.

### RNA expression studies (qPCR)

Total RNA was isolated by using the RNeasy Plus Mini kit (QIAGEN, Hilden, Germany) and quantified by using a Nanodrop 2000 UV visible spectrophotometer. One microgram of mRNA was reverse transcribed into cDNA by using an QuantiTect Reverse Transcription Kit (QIAGEN), according to the manufacturer’s instructions.

Real-time qPCR was performed on cDNA by using StepOnePlus™ Real-Time PCR System (Applied bioscience) and Takyon™ ROX SYBR 2X MasterMix dTTP blue (Eurogentec). For this test, predesigned primer pairs (**Supp. Table S2**) were purchased from Eurogentec. Fold changes in expression were calculated by the delta Ct method using RPLP0 (36B4) gene as an endogenous control for mRNA expression. All fold changes were expressed as normalized to the untreated control. Measurements were done in triplicate.

### Sprouting assay

Dry Cytodex-3 Beads (Cytiva, United Kingdom) were hydrated in PBS (Sigma-Aldrich, United States) and resuspended (30,000 beads/mL), autoclaved and stored in 4°C as per manufacturer’s instructions. Adequate e number of cells prepared and mixed with certain number of beads in the corresponding media of the cells. They will remain in incubator for 6 hours. Cell-coated beads were transferred to a 35mm petri dishes in 5 mL of complete media and left overnight. The day after, beads coated with cells are washed from the flasks, mixed with 2mg/ml bovine fibrinogen (Sigma-Aldrich) solution in PBS and distributed 2 ml/ well in 24-well plate. Thrombin (Sigma-Aldrich) is used as the enzyme to solidify fibrinogen. Normal human dermal fibroblasts were seeded onto each well at a concentration of 20,000 cells/well before plates were returned to a humidified incubator. The experiment terminated at 24 h post gel preparation by fixation (4% PFA) and then staining with 1:500 phalloidin at RT. Plates were imaged at x10 objective and images processed and quantified using ImageJ.

### Proteome Profiler

Cytokines present in the supernatant of LEC and HUVEC cells were quantified using the Proteome Profiler Human Chemokine Array kit (R&D Systems Cat. No. ARY017) and the human angiogenesis array kit (R&D Systems Cat. No. ARY007) according to manufacturer’s instructions. Briefly, the conditioned media samples 48 h post seeding from the cells were collected and applied on to the PVDF membranes that were precoated with the captured antibodies. The membranes were then washed with the washing buffer and incubated with the detection antibodies. The presence or absence of the cytokines were visualized with streptavidin-HRP mixture and the chemiluminescent HRP substrate. The immunoblot images were captured using film developer and the pixel density of the cytokine spots on the membranes were quantified using Image Lab software from Bio Rad.

### Immunofluorescence

Tumor tissues were fixed in PFA (4%) for half an hour immediately after dissection, then placed in a 30% sucrose solution overnight. Tumor tissues were frozen in OCT, then serially sectioned at five micrometers thick and allowed to sit at room temperature for half an hour to prepare for the immunostaining procedure. Prior to the application of the primary antibody, tissue sections were blocked for 60 minutes in phosphate-buffered saline (PBS) containing 1% bovine serum albumin (BSA), 5% horse serum, and 0.2% Triton. Incubation with primary antibodies was carried out either for one hour at room temperature or overnight at +4 °C. Negative controls were treated with blocking solution without the primary antibody. Incubation with fluorochrome-conjugated secondary antibodies (anti-rabbit AlexaFluor 488 and anti-rat AlexaFluor 647), specific to each primary antibody, was performed for 60 minutes at room temperature in the dark. DAPI was used to visualize nuclei.

### Transcriptome Analysis

#### RNA isolation and amplification

Total RNA was extracted from tissues lysed by TissueLyser II (QIAGEN) using phenol-chloroform extraction method with Trizol (Invitrogen) and Isopropanol (XILONG). The quality of the total RNA was controlled with concentration determination by using the Bioanalyzer 2100 (Agilent). All RNAs used in the present study had RNA Integrity Numbers (RINs) higher than 9.0.

#### RNA-Sequencing

RNA isolation, library preparation, and RNA-Seq were performed by BGI genomics (Hong Kong, China). Library preparation was performed from 1 µg of total RNA was used to prepare mRNA library using MGIEasy RNA Library Prep kit (MGI, China). Libraries were qualified on Bioanalyzer 2100 (Agilent) and quantified by PCR; single strand circle DNA (ssCir DNA) were formatted as the final library. The libraries of 9 samples were then amplified to make DNA nanoball (DNB) which and sequenced on DNBSEQ G400 sequencer (MGI, China) by pair-end 150, at least 30M clean reads was obtained for each library.

### Proteome Analysis

#### Migration on SDS-PAGE gel

Each sample was deposited on 10% acrylamide SDS-PAGE. After a short migration, gels were stained with colloidal blue overnight. After destaining, gels were scanned.

#### Identification of peptides/proteins by mass spectrometry

The bands of interest from the SDS-PAGE gel were cut and destained. Reduction disulfide bonds and alkylation of cysteine were performed, and proteins were finally digested overnight with trypsin. Resulting peptides were extracted from the gel, acidified and were injected on the Lumos with 146 min gradient OT/OT method. The spectra obtained were analyzed by Proteome Discoverer 3.1 with Sequest HT as the search algorithm, using a Human Uniprot database, including 2 missed cleavage sites by trypsin, carbamidomethylation of cysteine (+57 Da as static modification), oxidation of methionine (+16 Da as dynamic modification), acetylation of protein N-term (+42 Da as dynamic modification), Methionine loss (-131 Da as dynamic modification) and methionine loss/acetylation (+89 Da as dynamic modification).

#### Subcutaneous xenografts

This study was conducted in compliance with the National Charter on the ethics of animal experimentation. Our experiments were approved by the “Comité National Institutionnel d’Éthique pour l’Animal de Laboratoire (CIEPAL)” (reference: NCE/2023–823). Five million MDAMB231 cells (WT for one roundo f experiment and WT, Proton and Photon multil irradiated for another round) were injected, in medium containing 50% Corning® matrigel® matrix (VWR), subcutaneously, into the flank of 7-week-old NMRI-Foxn1nu/Foxn1nu female mice (Janvier Labs, Le Genest-Saint-Isle, France). Tumor volume was measured every other day with a caliper and calculated as follows:

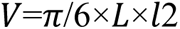

where *L* = tumor length; *l* = tumor width (in a 2D space, tangential to the mouse skin).

The experiment was stopped when the tumors of one experimental group reached a volume of 1000 mm^3^.

### Statistical analysis

Results are expressed as mean ± SD and analyzed using GraphPad Prism 9 (RRID:SCR_002798) software. The unpaired Student’s *t* test was used to determine the difference between 2 independent groups. For multiple comparison analyses, one-way ANOVA test with Turkey’s correction was used. Evaluation of the gaussian distribution of the data was performed prior to the *t* test or ANOVA. Normal distribution of the data and variance similarity were verified using GraphPad Prism. A *P* value less than 0.05 was considered statistically significant. All experiments were performed with a minimum of *N* = 3 biological replicates and *n* = 3 technical replicates. For *in vivo* studies, two-tailed Mann-Whitney U test was utilized to compare two independent groups and the sample size was determined using the methods described by Berndtson *et al*. [18]. Predetermined exclusion criteria included the absence of signal at the start of the experiment.

## Results

### The sensitivity of cells within the tumor microenvironment to conventional X irradiation surpasses that of P irradiation while they secret VEGFC to the same level

The primary objective of this study was to assess the risk of relapse in TNBC patients following surgery, specifically focusing on adjuvant radiotherapy with either X or P. To achieve this, we aimed to elucidate the distinct effects of these radiation types on various components of the tumor microenvironment, including vascular endothelial cells (VEC; TERT-immortalized microvascular endothelial cells (TIME) and primary Human Umbilical Vein Endothelial Cells (HUVEC)), lymphatic endothelial cells (LEC) and human fibroblasts (FHN). These cell types were selected based on their anticipated differential production of vascular endothelial growth factors, pivotal for the development of blood and lymphatic networks, representing key routes for metastatic dissemination. Except for FHN cells, which were exposed to 8 Gray (Gy), all other cell types were subjected to escalating radiation doses to evaluate their relative sensitivity. Notably, a dose-dependent increase in cell mortality was observed across TIME, HUVEC and LEC cells 48 hours post-irradiation (Figure S1A-F), from 2 Gy to 8 Gy. To highlight the overall biological effects, cells were exposed to either a single or multiple irradiations at the highest dose of 8 Gy.

In evaluating the relative sensitivity of different tumor microenvironment cell types, LEC, VEC and FHN, to X or P radiation, we employed clonogenic assays to encompass both acute and long-term killing effects (**Figure 1A**). Contrary to conventional beliefs, our observations revealed a noteworthy distinction in clonogenic survival following a single high-dose irradiation of 8 Gy, with a higher number of remaining colonies after P irradiation compared to X irradiation. This suggests inherent differences in how each type of irradiation influences the tumor microenvironment, potentially creating conditions favorable for relapse through blood or lymphatic vessels. This finding aligns with the importance of sentinel lymph node positivity, not just as a diagnostic marker but also as a prognostic indicator. Given the critical role of lymphangiogenesis in cancer metastasis and our previous findings in head and neck as well as renal tumors [16, 19], where elevated VEGFC expression was identified as a stress response factor, we hypothesized that VEGFC could be a primary driver of TNBC relapse and metastasis to the axillary lymph node following irradiation.

**Figure 1.**
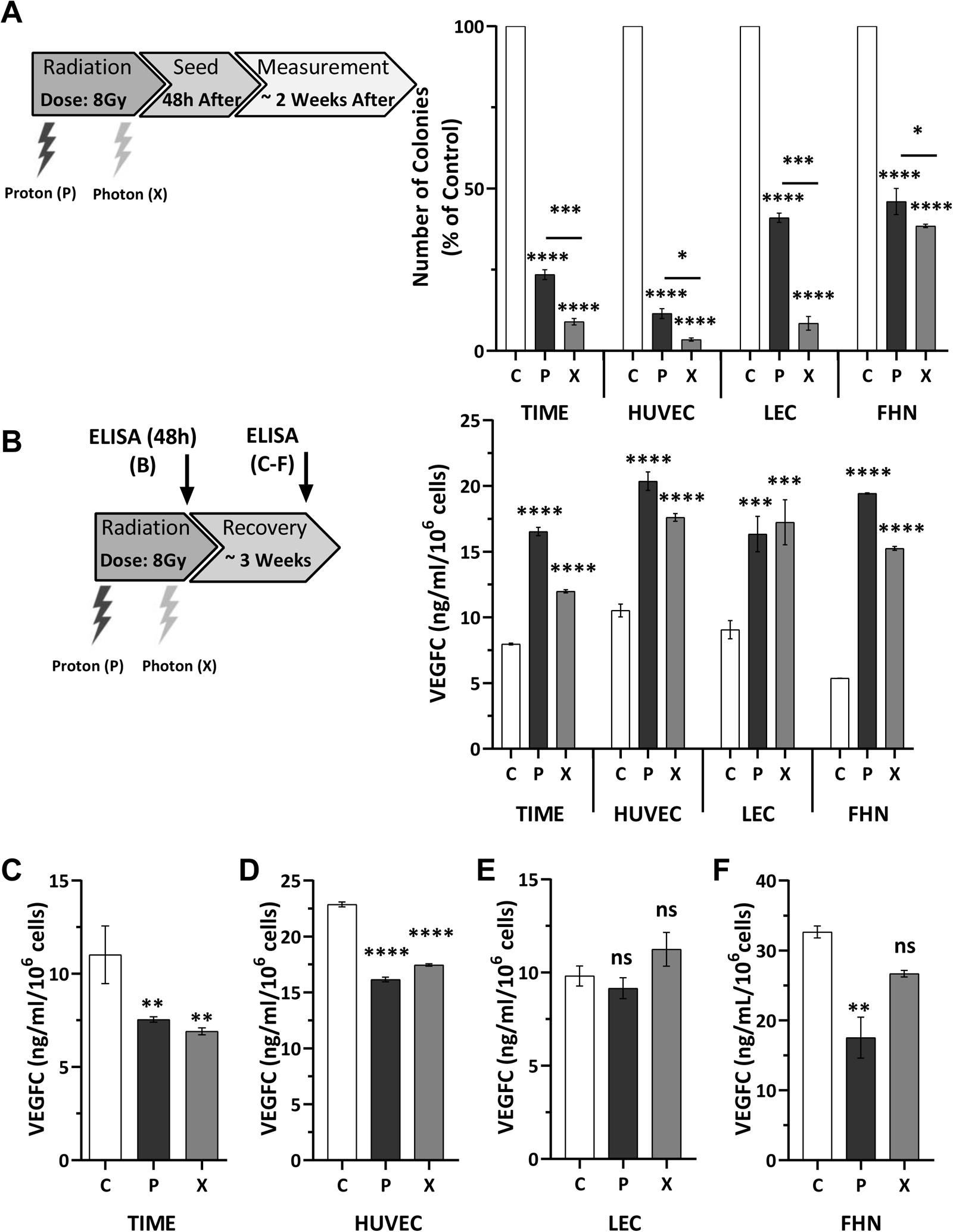
Sensitivity of cells from the tumor microenvironment to proton (P) and photon (X) irradiation, along with the subsequent increased secretion of VEGFC in these cells post-irradiation. **(A)** The clonogenic potential of cells from the tumor microenvironment (TIME, HUVEC, LEC and FHN) was evaluated following irradiation with P and X beams at a dose of 8Gy. Clonogenic assays were performed by seeding cells 48 hours post-radiotherapy and assessing colony formation over a two-week period. **(B)** Measurement of VEGFC protein levels secreted in the supernatant of normal cells from the tumor microenvironment (TIME, HUVEC, LEC, FHN) 48 hours after a single round of P and X irradiation (8Gy), compared to their respective controls (C), using ELISA. **(C-F)** Assessment of VEGFC protein levels in the supernatant of TIME, HUVEC, LEC and FHN cells two weeks post single round of P and X irradiation (8Gy), compared to their respective controls (C). The results are presented as the mean of at least three independent experiments ± standard deviation (SD). Statistical analysis was performed using one-way ANOVA to compare differences between the control and irradiated groups (P or X). Statistical significance is denoted as follows: *P < 0.05, **P < 0.01, ***P < 0.001, ****P < 0.0001, NS (non-significant).

To investigate this hypothesis, we assessed the expression of VEGFC in various cell types from the tumor microenvironment: LEC, HUVEC, TIME and FHN. VEGFC expression was measured 48 hours post irradiation with either P or X radiation. In concordance with the dose-dependent effect observed on cell survival, a dose-dependent increase in VEGFC expression was noted in TIME, HUVEC and LEC cells with both P (Figure S2A-C) and X irradiation (Figure S2D-F) from 2 to a maximum dose of 8 Gy. So, we observed an increase in VEGFC secretion in all these cells following both P and X irradiation at 8 Gy (**Figure 1B**).

To evaluate the sustainability of this response and thus the enduring role of VEGFC, we assessed its expression later after irradiation (3-4 weeks post irradiation) in TIME, HUVEC, LEC, and FHN. The results underscored the transient nature of the effect, as VEGFC levels decreased compared to the control, indicative of an intense stress response followed by a subsequent decline after the stressor ceased (**Figure 1C-F**).

These findings strongly suggest that VEGFC is a key player in the adaptive response to the substantial stress induced by radiation, whether with X or P.

Subsequently, treatments tailored to specific tumor types may further modulate tumor cells through their genetic plasticity. Moreover, cells within the tumor microenvironment, despite possessing lower genetic plasticity, may undergo genetic modifications contributing to adaptive responses, either transiently or chronically, if they withstand the aforementioned treatments.

These findings underscore the complex interplay between tumor cells, treatment modalities, and the microenvironment.

### Repeated exposure of vascular endothelial cells to multiple rounds of irradiation results in the acquisition of profibrotic properties

To simulate the impact of radiation treatments administered to cancer patients on individual cells within the tumor microenvironment, we subjected cells to a multi-irradiation process, similar to the multiple radiation sessions undergone by patients. Our objective was to elucidate the potential deleterious effects induced by such repetitive radiation exposures on both normal and residual tumor cells following surgical intervention. We conceptualized radiation as a stressor capable of inducing genetic adaptations that lead to the production of detrimental cytokines contributing to local or distant metastasis.

To investigate this hypothesis, TIME cells, integral to the formation of blood vessels crucial for metastatic spread, underwent several rounds of irradiation by either X or P. Remarkably, after multiple rounds, the percentage of remaining cells approached 100% (**Figure 2A**). Subsequent analyses revealed that multi-irradiated TIME cells exhibited heightened proliferative capabilities, particularly pronounced in P-adapted cells (**Figure 2B**). Moreover, these adapted cells demonstrated an increased migratory capacity in Transwell assay (**Figure 2C**). Initially interpreted as augmented angiogenic properties, further investigation through sprouting assays contradicted this expectation, revealing a substantial impairment in the ability to form pseudo blood vessels in a 3D model, as evidenced by reduced length and number of sprouts (**Figure 2D-F**). Contrary to the anticipated increase in angiogenesis, our findings suggested that reduced sprouting of multi-irradiated endothelial cells indicates the induction of fibrosis through an endothelial-to-mesenchymal transition (Endo-MT) as it previously observed in lung cancer [20]. To substantiate this hypothesis, we examined the expression of several pro-fibrotic genes using qPCR. Notably, transforming growth factor beta (TGFβ), fibrillin 1 (FBN1), heparan sulfate proteoglycan 2 (HSPG2), Von Willebrand factor (vWF), fibroblast activation protein (FAP) and α-smooth muscle actin (αSMA) were almost all upregulated in radiation-adapted cells (**Figure 2G**).

**Figure 2.**
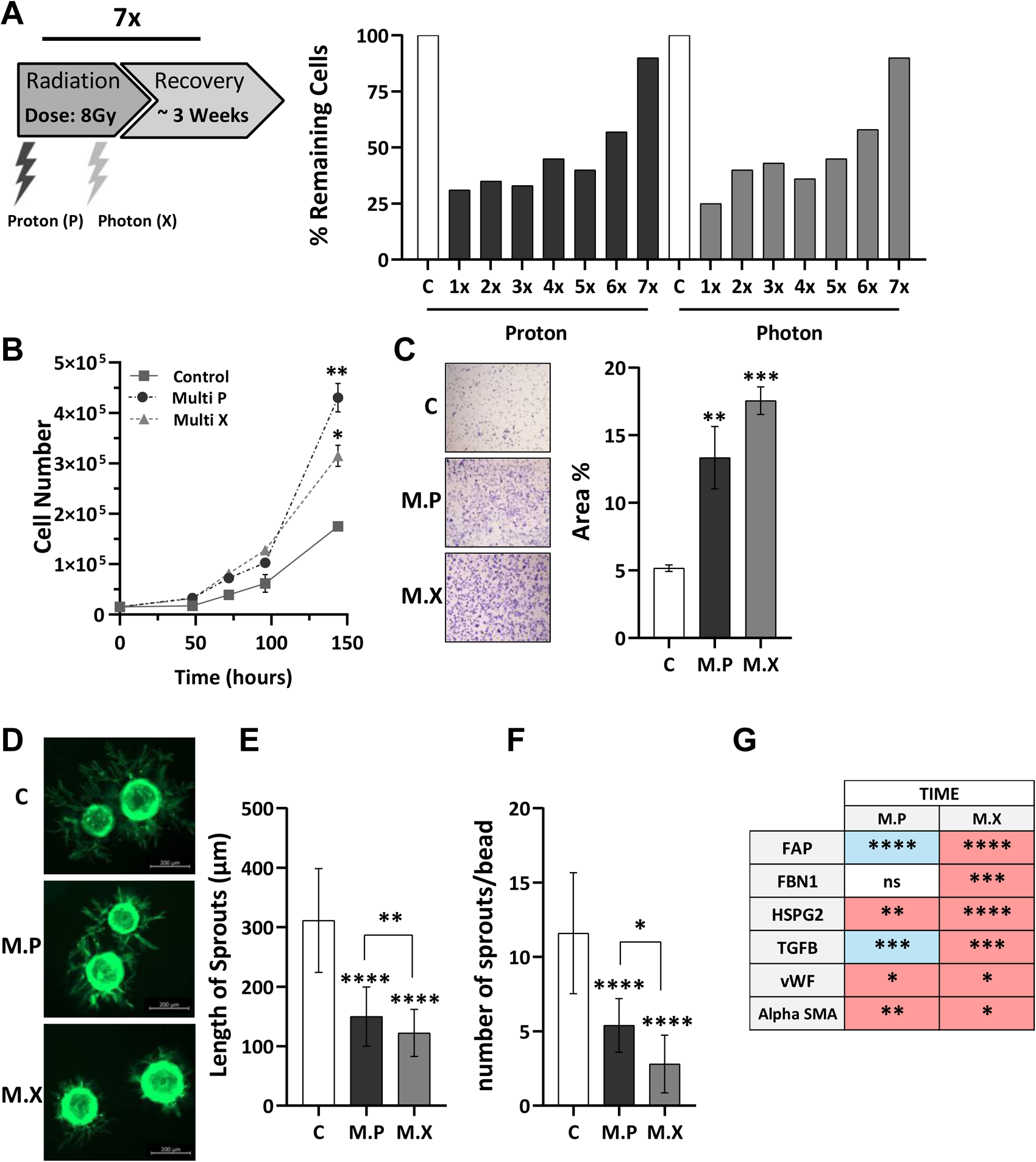
Irradiation of vascular endothelial cells induces a pro-fibroblastic phenotype. **(A)** Assessment of remaining TIME cells 48 hours following seven rounds of P and X irradiations (dose: 8 Gy) termed as multi-irradiated TIME cells, compared to their respective controls (C, 0 Gy). **(B)** Analysis of the proliferation rate (cell counts) of multi-irradiated TIME cells with P and X irradiation over a 144-hour period, compared to their respective control. **(C)** Evaluation of the migration capacity (Transwell assays) of multi-irradiated TIME cells with P and X irradiation, compared to the corresponding controls (C). **(D-F)** Assessment of the sprouting ability of multi-irradiated TIME cells with P and X irradiation, compared to the corresponding controls (C) through 3D sprouting on Cytodex beads. **(G)** Quantitative gene expression analysis of multi-irradiated TIME cells with P and X irradiation, compared to the corresponding Control (C). Red indicates upregulation of the gene compared to the respective control, while blue represents downregulation. The results are presented as the mean of at least three independent experiments ± SD. Statistical analysis was performed using one-way ANOVA to compare differences between the control and irradiated groups (P or X). Statistical significance is denoted as follows: *P < 0.05, **P < 0.01, ***P < 0.001, ****P < 0.0001, NS (non-significant).

These results strongly suggest that radiotherapy amplifies a profibrotic phenomenon in endothelial cells that may persist post-surgery at the scar site. This fibrosis, reminiscent of descriptions in various tumors, including BC, may foster a pro-tumor environment through the secretion of several proinflammatory, pro-angiogenic, and pro-lymphangiogenic cytokines and growth factors [21–23].

Moreover, the altered phenotype of multi-irradiated TIME cells suggests the involvement of other factors in addition to VEGFC. Their expression may be stimulated through the VEGFC-dependent signaling pathway, potentially amplifying the initial impact mediated by VEGFC on blood and lymphatic vessel networks.

### The transient induction of VEGFC following irradiation with X and P is concomitant or followed by the activation of other pro-angiogenic and pro-lymphangiogenic cytokines

To identify the factors involved in the long-term response initiated by the initial surge of VEGFC, we conducted a cytokine array analysis using conditioned media from HUVEC and LEC (Figure S3A-B). While this approach provides a broad overview of the pro-angiogenic and lymphangiogenic secretome modifications, it may not capture the entirety of the global secretome. The analysis revealed common factors as well as differences in the secretome shaped by P and X irradiation, potentially influencing the behavior of blood and lymphatic vessels post-irradiation. Among the 55 evaluated factors, some were not changes, some were modified in the same way in response to X and P irradiations and were differentially expressed in P and X-irradiated; five were differentially expressed in LEC (Activin, artemin, DPPIV, endostatin, and VEGFC) (Figure S3C) and 7 in HUVEC cells (Activin, angiopoietin 2, artemin, endostatin, IGFBP2, PDGFA, and VEGFC) (Figure S3D). Notably, four factors (Activin, artemin and VEGFC) were co-regulated by both types of irradiations.

In HUVEC, only P irradiation stimulated the expression of detrimental factors (angiopoietin 2 and IGFBP2), along with the controversial factor endostatin, which has been reported to exhibit both pro-and anti-tumoral effects. Importantly, the decrease in the pro-tumor factor artemin was observed only for X irradiation. A similar decrease in artemin was also noted in LEC, but only in response to X irradiation. However, the detrimental factor DPPIV exhibited an increase with X irradiation and a decrease with P irradiation (Figure S3C-D). Overexpression of all these genes, included VEGFC, correlates with a shorter overall cancer survival rate (Figure S4).

These results suggest distinct adaptation pathways to X and P irradiation, as well as potential crosstalk between different factors whose expression is stimulated by irradiation. Notably, previous studies have suggested crosstalk between IGFBP2, as well as between angiopoietin 2 and VEGFC [24].

### Multi-irradiation of TNBC cells promotes the development of a pro-metastatic phenotype

To evaluate the sensitivity of TNBC cells to P and X radiations, we initially irradiated a variety of TNBC cell lines, including human (MDAMB231, BT549, CAL51, and EO) and murine (4T1) models. Using clonogenic assays to assess both acute and long-term killing effects, we observed a significant difference in clonogenic survival following a single high-dose irradiation of 8 Gy, with more colonies remaining after P irradiation compared to X irradiation. This trend was consistent across most cell lines, except for BT549 cells, where an equivalent number of colonies persisted post-irradiation despite the radiation type (**Figure 3A**).

**Figure 3.**
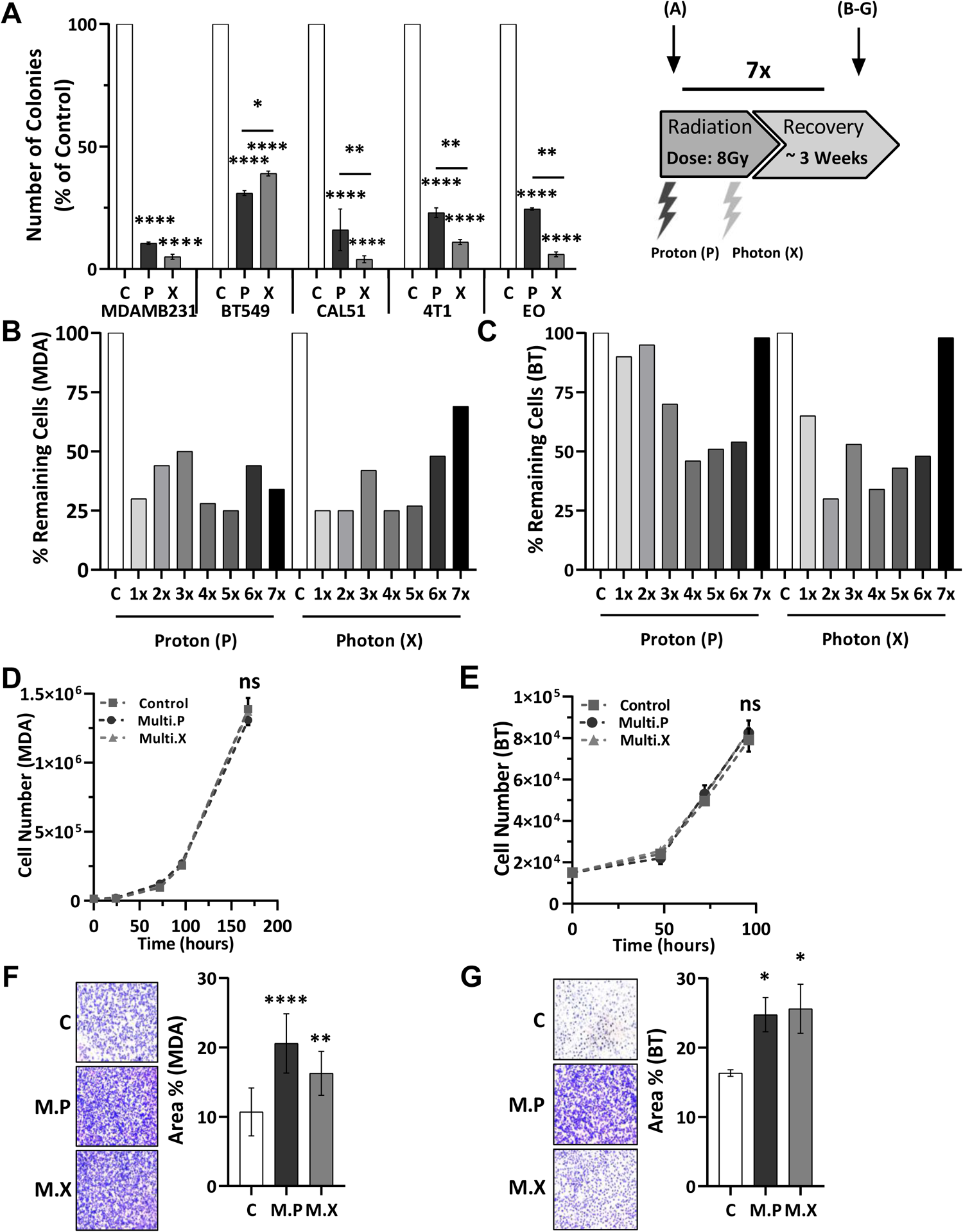
Multi-irradiated TNBC cells acquired pro-metastatic properties. **(A)** The clonogenic potential of breast tumor cells (MDAMB231, BT549, CAL51 for human cells; 4T1, and EO for mouse cells) was evaluated following irradiation with P and X beams at a dose of 8Gy. Clonogenic assays were performed by seeding cells 48 hours post-radiotherapy and assessing colony formation over a two-week period. **(B-C)** Assessment of remaining MDAMB231 and BT549 cells 48 hours following seven rounds of P and X irradiations (dose: 8 Gy) termed as multi-irradiated BT549 and MDAMB231 cells, compared to their respective controls (C, 0 Gy). **(D-E)** The proliferation rate (cell counts) of MDAMB231 and BT549 cells was evaluated following seven rounds of P and X irradiations (8 Gy), referred to as multi-irradiated cells, compared to their respective controls. **(F-G)** The migration ability of multi-irradiated (P and X) BT549 and MDAMB231 was assessed compared to their corresponding controls. The results are presented as the mean of at least three independent experiments ± SD. Statistical analysis was performed using one-way ANOVA to compare differences between the control and irradiated groups (P or X). Statistical significance is denoted as follows: *P < 0.05, **P < 0.01, ***P < 0.001, ****P < 0.0001, NS (non-significant).

To better elucidate the involvement of radiotherapy treatment of TNBC cells, we generated cell populations resilient to repeated exposure to X and P irradiation. This investigation encompassed two predominant TNBC models, MDAMB231 and BT549. MDAMB231 cells exhibited high sensitivity, with only 25% of cells persisting 48h after each X or P irradiation (**Figure 3B**). Conversely, BT549 cells displayed inherent resistance to irradiation, with 90% of cells remaining viable 48 hours post-irradiation, irrespective of the radiation type (P or X) (**Figure 3C**). Notably, even with a gradual decrease in cell count over successive irradiation rounds, BT549 cells demonstrated substantial resilience after seven cycles of irradiation. Although a slight enrichment of cell numbers was observed following the seventh round, these cells were deemed adapted to irradiation.

Upon selecting these adapted cells, we assessed various hallmarks of aggressiveness, including proliferation and migration capabilities. Proliferation rates of adapted MDAMB231 and BT549 cells exposed to either X or P radiation remained comparable to control cells (**Figure 3D-E**). However, migration abilities assessed through Transwell assays showed a significant increase for both MDAMB231 and BT549 cells (**Figure 3F-G**), consistent with observations in X-resistant medulloblastoma cells [25]. This finding parallels the observations in adapted TIME endothelial cells (**Figure 2C**).

Consequently, multi-irradiations induce modifications in the pro-invasive phenotype characteristic of highly invasive cells, that can lead to increase the propensity for metastatic formation for the remaining cells post radiotherapy.

### VEGFC emerges as a pivotal regulator in the post-irradiation behavior of TNBC

Given the robust induction of VEGFC following various cancer treatments, including radiotherapy, across diverse cancer models [16, 19, 25], we explored the impact of X and P irradiation on VEGFC expression in two distinct human TNBC cell lines (MDAMB231 and BT549 characterized by high basal VEGFC levels (Figure S5B). Both X and P irradiations markedly elevated VEGFC mRNA (**Figure 4A**) and protein (**Figure 4B**) levels in these cell lines, consistent with previous observations in head and neck tumor cells [16]. To further explore the genetic mechanisms associated with enhanced angiogenesis and lymphangiogenesis following multiple irradiations that simulate clinical treatments, we evaluated VEGFC levels alongside other relevant genes using cytokine array analysis. Notably, both VEGFC mRNA and protein levels were consistently upregulated in adapted MDAMB231 and BT549 cells after exposure to either P or X irradiation. While the increase in VEGFC protein was significantly pronounced in MDAMB231 cells, BT549 cells showed only a trend toward increased protein levels (**Figure 4C-D**). As shown in the tumor microenvironment, normal cells exhibited higher VEGFC expression compared to tumor cells (**Figure 1B and 4A**), including two TNBC cell lines of human and murine origin that lacked VEGFC expression (Figure S5B). To explore the autocrine proliferative potential of VEGFC, like our previous findings in head and neck and renal tumor cells [16, 19], we assessed the expression of VEGFC pathway components. This included VEGFC itself and its receptors (VEGFR2 and VEGFR3), as well as co-receptors neuropilin 1 (NRP1), neuropilin 2 (NRP2), and CD146. Strikingly, not all available human TNBC cell lines expressed all members of the VEGFC pathway, and some, such as CAL51, did not express VEGFC (DU4475, HCC1599, BT20, MDAMB468, (Figure S5A). Considering the autocrine and paracrine effects of VEGFC, we investigated the relationship between VEGFC expression and survival using available databases and KM plotter software. High VEGFC expression in TNBC correlated with shorter survival (**Figure 4E**), suggesting that basal VEGFC expression or treatment-induced VEGFC induction, as observed in head and neck or renal tumors, could lead to a poorer clinical outcome. These findings underscore the importance of considering the distinct impact of radiation types on the tumor microenvironment and highlight potential implications for treatment strategies in TNBC patients. Additionally, artemin, VEGFA and FAP mRNA levels exhibited heightened expression in both cell lines post-irradiation with P and X evaluating by qPCR (**Figure 4F**). All these factors correlated with lower survival in TNBC (Figure S4). Intriguingly, the mRNA levels of certain genes such as angiopoietin 2, IGFBP2, and TGFβ showed variable induction across different conditions and cell lines (**Figure 4F**). These results suggest that the genetic alterations of tumor cells may vary subtly depending on the irradiation regimen, potentially influencing the tumor microenvironment and contributing to the risk of future disease relapse

**Figure 4.**
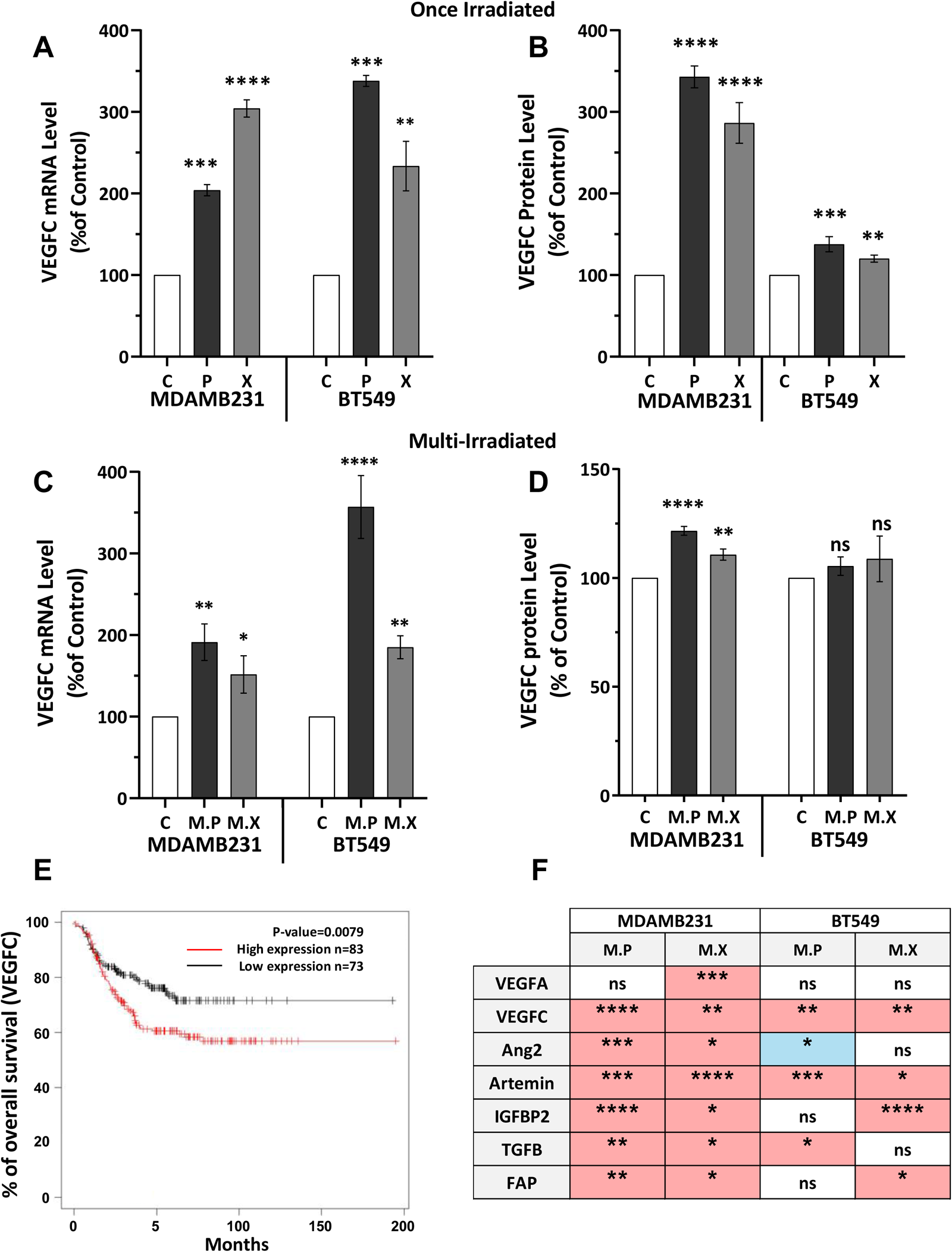
Single or multiple irradiations stimulate VEGFC production by TNBC cells and initiate a pro-angiogenic/pro-fibroblastic phenotype. **(A)** Measurement of VEGFC mRNA levels in MDAMB231 and BT549 cells 48 hours after a single round of P or X irradiations (8 Gy), referred to as single irradiated, compared to their corresponding controls (C) using quantitative PCR. **(B)** Quantification of secreted VEGFC protein levels in supernatant of single P or X irradiated MDAMB231 and BT549 cells 48 hours post irradiation compared to respective controls (C) using ELISA. **(C)** Evaluation of VEGFC mRNA levels of MDAMB231 and BT549 cells after seven rounds of P or X irradiations, referred to as Multi Proton (M.P) and Multi Photon (M.X), respectively, compared to their corresponding controls (C). **(D)** Quantification of secreted VEGFC protein levels in the supernatant of M.P and M.X irradiated of MDAMB231 and BT549 cells compared to their respective controls (C) using ELISA. **(E)** Kaplan-Meier analysis of overall survival of TNBC patients using the Kaplan Meier softaware (https://kmplot.com/analysis/). OS was calculated from patient subgroups with mRNA levels of VEGFC that were less or greater than the best cut-off value. **(F)** Quantitative gene expression analysis of M.P and M.X MDAMB231 and BT549 compared to the corresponding Control (C). Red indicates upregulation of the gene compared to the respective control, while blue represents downregulation. The results are presented as the mean of at least three independent experiments ± SD. Statistical analysis was performed using one-way ANOVA to compare differences between the control and irradiated groups (P or X). Statistical significance is denoted as follows: *P < 0.05, **P < 0.01, ***P < 0.001, ****P < 0.0001, NS (non-significant).

### The VEGFC gene knock-out significantly enhances the sensitivity of TNBC cells to irradiation

Recognizing the pivotal role of VEGFC in conferring resistance to radiation, we utilized CRISPR/Cas9 technology to knock-out its gene, a methodology previously employed in other models [25, 26]. Consequently, independent control and VEGFC-/-clones of MDAMB231 and BT549 cells were subjected to either X or P irradiation. Remarkably, while approximately 50% of cells remained viable 96-168 hours after irradiation in the control group, VEGFC-/-cells exhibited a drastic reduction in cell count, with almost no viable cells remaining both for MDAMB231(**Figure 5A-B**) and BT549 cells (**Figure 5C-D**).

**Figure 5.**
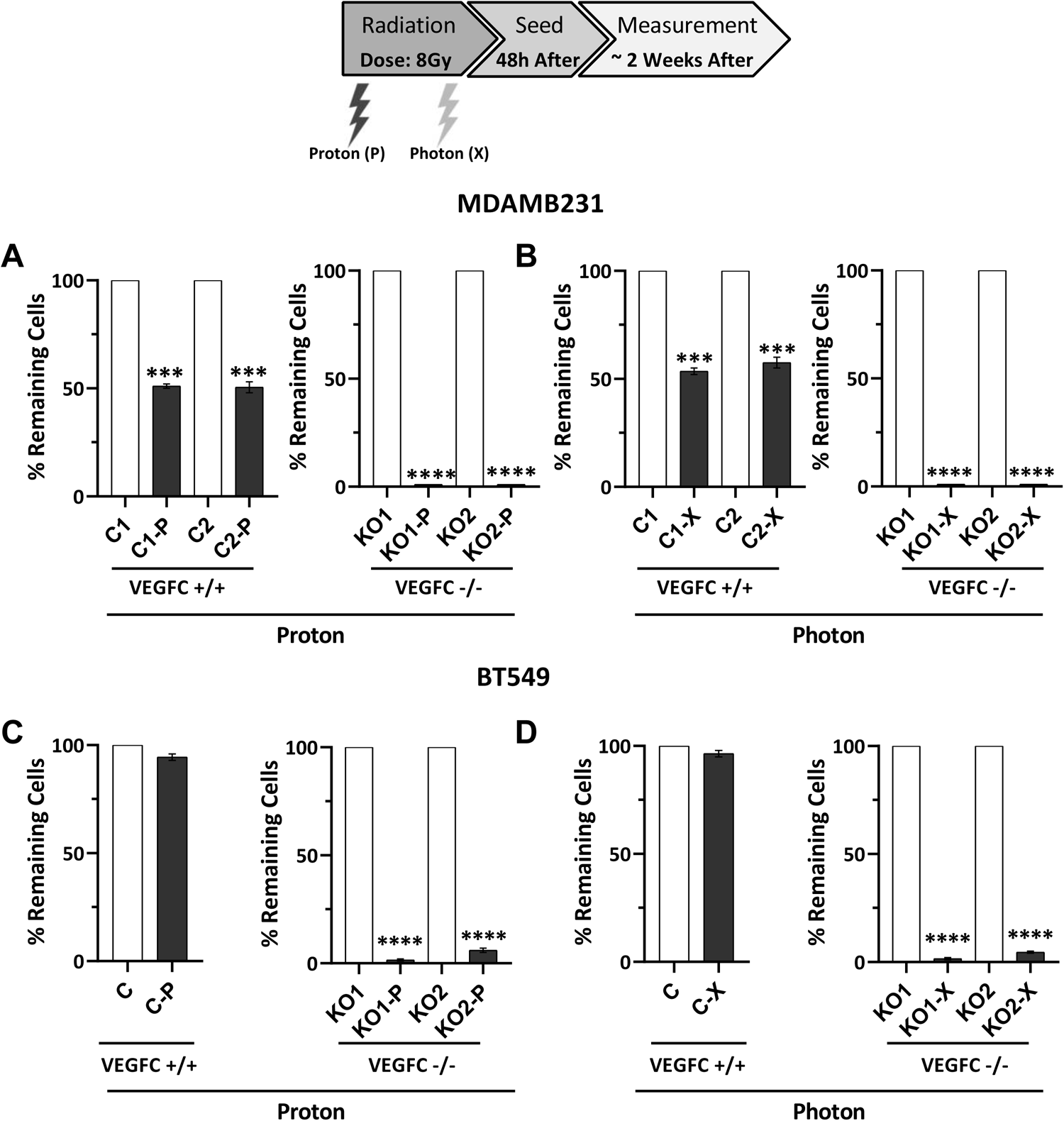
VEGFC knock-out cells are more sensitive to irradiation. **(A-B)** Cell counts were performed on CRISPR/Cas9-edited MDAMB231 cells was conducted with empty vector serving as the Control (C) and VEGFC knockout (KO) as the experimental condition two weeks after exposure to P or X irradiation (8 Gy). Two clones were assessed for each condition. **(C-D)** Cell count was performed on CRISPR/Cas9-edited BT549 cells, with the empty vector serving as the control (C) and VEGFC knockout (KO) as the experimental condition. Two weeks after exposure to P or X irradiation (8 Gy). One clone from the control group and two clones from the KO group were assessed for each condition. The results are presented as the mean of at least three independent experiments ± SD. Statistical analysis was performed using one-way ANOVA to compare differences between the control and irradiated groups (P or X). Statistical significance is denoted as follows: *P < 0.05, **P < 0.01, ***P < 0.001, ****P < 0.0001, NS (non-significant).

To ascertain whether this heightened sensitivity to radiotherapy translates to other treatment modalities, we subjected both control and VEGFC-/-MDAMB231 clones to chemotherapeutic agents commonly employed in TNBC treatment regimens, such as paclitaxel (Taxol) (Figure S6A) and doxorubicin (Figure S6B-C). VEGFC-/-KO cells demonstrated either comparable sensitivity or, depending on the dosage, even exhibited greater resistance compared to control cells. This observation implies that VEGFC may exert a more specific influence on the sensitivity to radiation within these TNBC models.

### Deletion of the VEGFC gene enhances migration capabilities

To investigate whether the knock-out of VEGFC could alter specific markers of aggressiveness, we assessed the relative proliferation and migration abilities in MDAMB231 and BT549. Two independent wild-type and VEGFC-/-clones were subjected to testing. The clonogenic assays (**Figure 6A-B**) and proliferation rate, evaluated through cell counting (**Figure 6C-D**), remained comparable between control and VEGFC-/-cells in both cell lines. However, VEGFC-/-in MDAMB231 and BT549 cells showed higher migration capacity, assessed via Traswell assays (**Figure 6E-F**). This observed increase in migration ability aligns with findings from other models, including kidney tumors and medulloblastoma cells [25, 26]. Thus, across independent tumor models, VEGFC disruption appears to promote a pro-migration phenotype, a characteristic associated with heightened metastatic potential.

**Figure 6.**
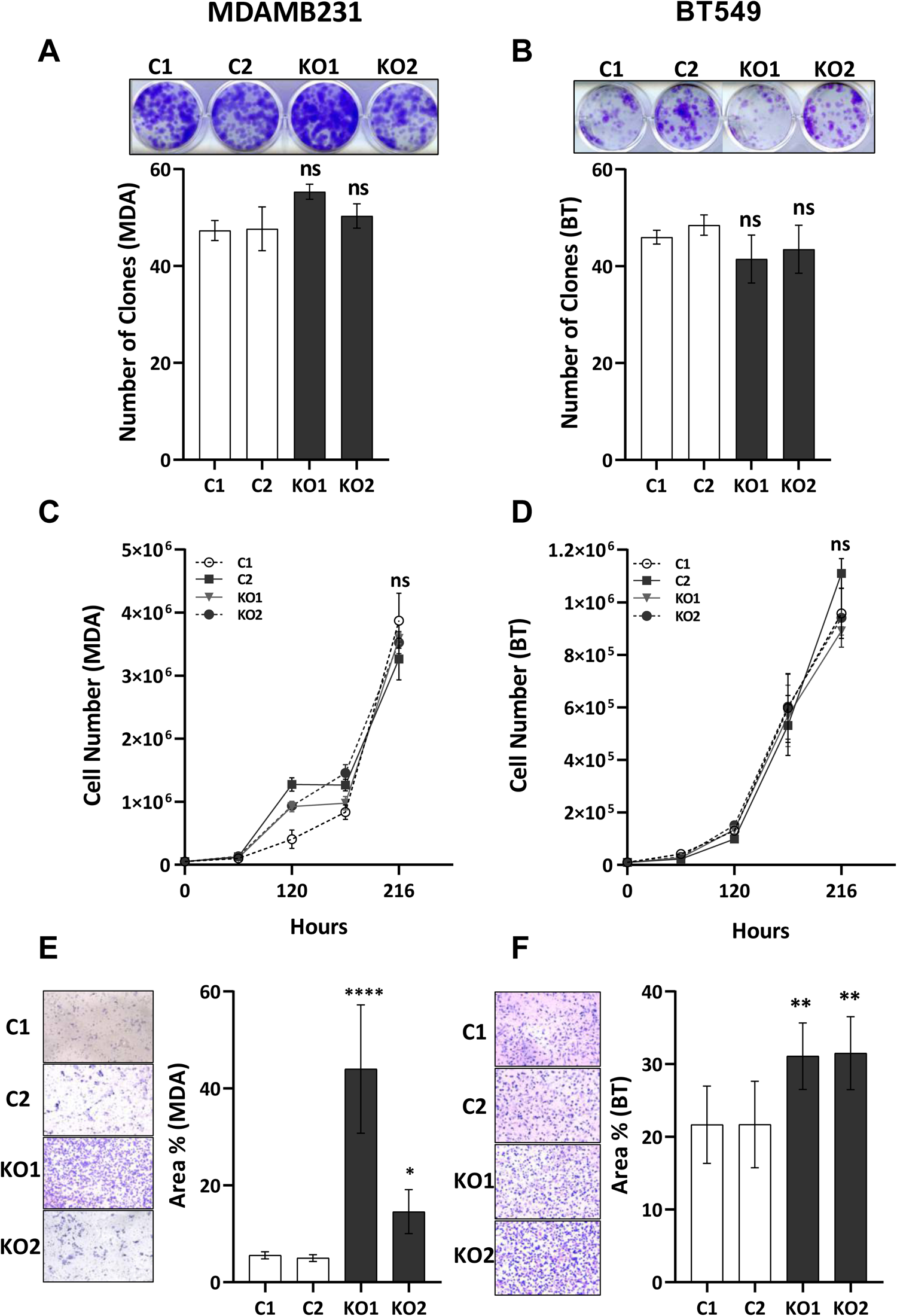
VEGFC knock-out cells presents with increased migration abilities. **(A-B)** Assessment of the clonogenic ability of the same CRISPR/Cas9-edited MDAMB231 and BT549 cells during two weeks. Two clones were evaluated for each condition. **(C-D)** Cell counts of CRISPR/Cas9-edited MDAMB231 and BT549 cells, including control (C) and knockout (KO) variants, were monitored over a period of 216 hours. Two clones were evaluated for each condition. **(E-F)** The migration capacity of the same CRISPR/Cas9-edited MDAMB231 and BT549 cells, measured by transwell assay. Three clones were evaluated for each condition The results are presented as the mean of at least three independent experiments ± SD. Statistical analysis was performed using one-way ANOVA to compare differences between the control and irradiated groups (P or X). Statistical significance is denoted as follows: *P < 0.05, **P < 0.01, ***P < 0.001, ****P < 0.0001, NS (non-significant).

### TNBC cells adapted to multi-irradiations by P generate bigger tumors but less aggressive than X tumors

Based on the aggressive behavior observed *in vitro* in TNBC cells adapted to multi-irradiation, along with our previous findings in head and neck tumors demonstrating increased *in vivo* aggressiveness of cells adapted to multi-irradiations [16], we conducted experiments to assess the tumorigenic capacity of TNBC P and X-adapted cells in nude mice. Tumors generated with P-adapted cells (P tumors) exhibited a more rapid onset compared to tumors generated with control or X-adapted (X tumors) cells (**Figure 7A**). The incidence of tumor formation (i.e., presence of tumors in mice) reached 100% for P tumors, while it was approximately 60% for the X tumors and 40% for controls. Furthermore, tumor volumes were notably larger for P tumors, displaying exponential growth, whereas X tumors, though lower, still exhibited larger volumes than control tumors (**Figure 7C**). At the end of the experiment, both tumor volume and weight were significantly higher, particularly for P tumors (**Figure 7C-E**). These findings suggest that TNBC cells that have survived multiple irradiations, especially by P, possess the capability to generate much more aggressive tumors.

**Figure 7.**
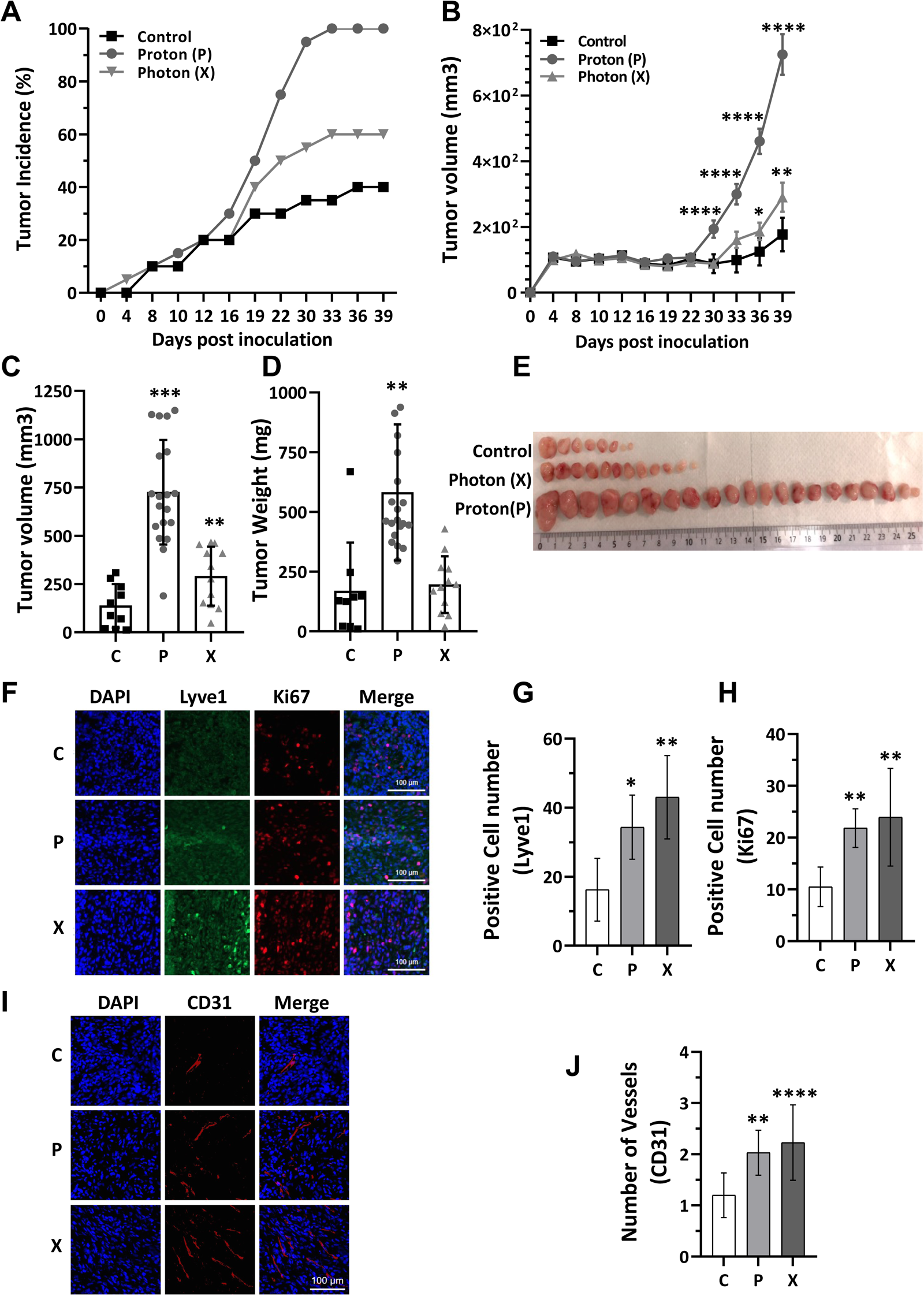
TNBC cells multi-irradiated by P generate bigger tumors than cells multi-irradiated by X. The evaluation of tumors generated following xenografting of either non-irradiated (Control), seven rounds of P irradiated (8 Gy), or X irradiated (8 Gy) MDAMB231 cells in immunodeficient mice. **(A)** Comparison of tumor incidence between the control, P, and X groups under the same conditions. **(B)** Monitoring of tumor growth in the three groups, including control and irradiated groups, over a period of 39 days. **(C)** Measurement of tumor volumes on the last day of the experiment measured for all three groups. **(D)** Measurement of tumor weights on the last day of the experiment measured for all three groups. **(E)** Representative image of the tumor xenografts showcasing the morphology and growth of the tumors in each experimental group. Immunofluorescence (IF) of lymphatic, proliferative and vascular markers in murine xenografts. **(F-H)** Representative images of LYVE1 (lymphatic endothelial cells, green)/Hoechst (nuclei, blue) and Ki67 (proliferative cells, red)/Hoechst (nuclei, blue) staining, showing different patterns of lymphatic vessels development in P, X and C tumor groups **(I-J)** Representative images CD31 (endothelial cells, red)/Hoechst (nuclei, blue) staining, showing anarchic blood vessels structures in P, X and C tumor groups Differences between the control and irradiated groups (P or X) were analyzed using one-way ANOVA. Statistical significance was denoted as follows: *P < 0.05, **P < 0.01, ***P < 0.001, ****P < 0.0001, NS (non-significant).

To further explore the characteristics of P and X tumors, we investigated lymphatic, proliferative, and vascular markers—specifically Lyve1 (Lymphatic Vessel Endothelial Hyaluronan Receptor 1), Ki67, and CD31—using tumor tissue sections post-dissection. Surprisingly, we found higher expression of these markers in X tumors compared to P tumors, and both were significantly elevated compared to corresponding controls (**Figure 7F-J**).

To have deeper insight in the genetic modifications of P and X tumors, as we had human tumor cells inside a murine model, we did transcriptomic analysis of both human and mouse. We found revealed that X tumors exhibit a higher activated signature of angiogenesis and lymphangiogenesis compared to P tumors (**Figure 8A**). Concurrently, Gene Set Enrichment Analysis (GSEA) between X and P tumors showed pronounced expression of lymphangiogenesis and proliferation at the mouse level (**Figure 8B**). Consistent with these findings in mice, GSEA enrichment analysis at the human level identified biological pathways significantly over-represented in the list of differentially expressed genes, including angiogenesis, epithelial-mesenchymal transition (EMT), cancer module 516 (FGF1, ANGPT1, MMP19, JAG1, IL18, NRG1), and cancer module 138 (SMAD3, ETS1, VDR, RGS4, ENG, DNTT). These results indicate a higher aggressiveness of X tumors compared to P tumors (**Figure 8C**). These observations suggest that despite X tumors having a smaller volume, they exhibit greater molecular aggressiveness compared to P tumors. This is evidenced by increased proliferation (Ki67) and the presence of lymphatic (Lyve1) and vascular markers (CD31) (**Figure 7F-J**), which facilitate tumor dissemination. The higher levels of lymphatic markers in X tumors in addition to the result of transcriptomic analysis that shows higher lymphangiogenic profile in X tumors (**Figure 8A**), suggest the potential benefit of combining *in vivo* tumor growth studies with anti-VEGFC antibody treatment to validate the role of lymphangiogenesis in the prognosis of irradiated tumors.

**Figure 8.**
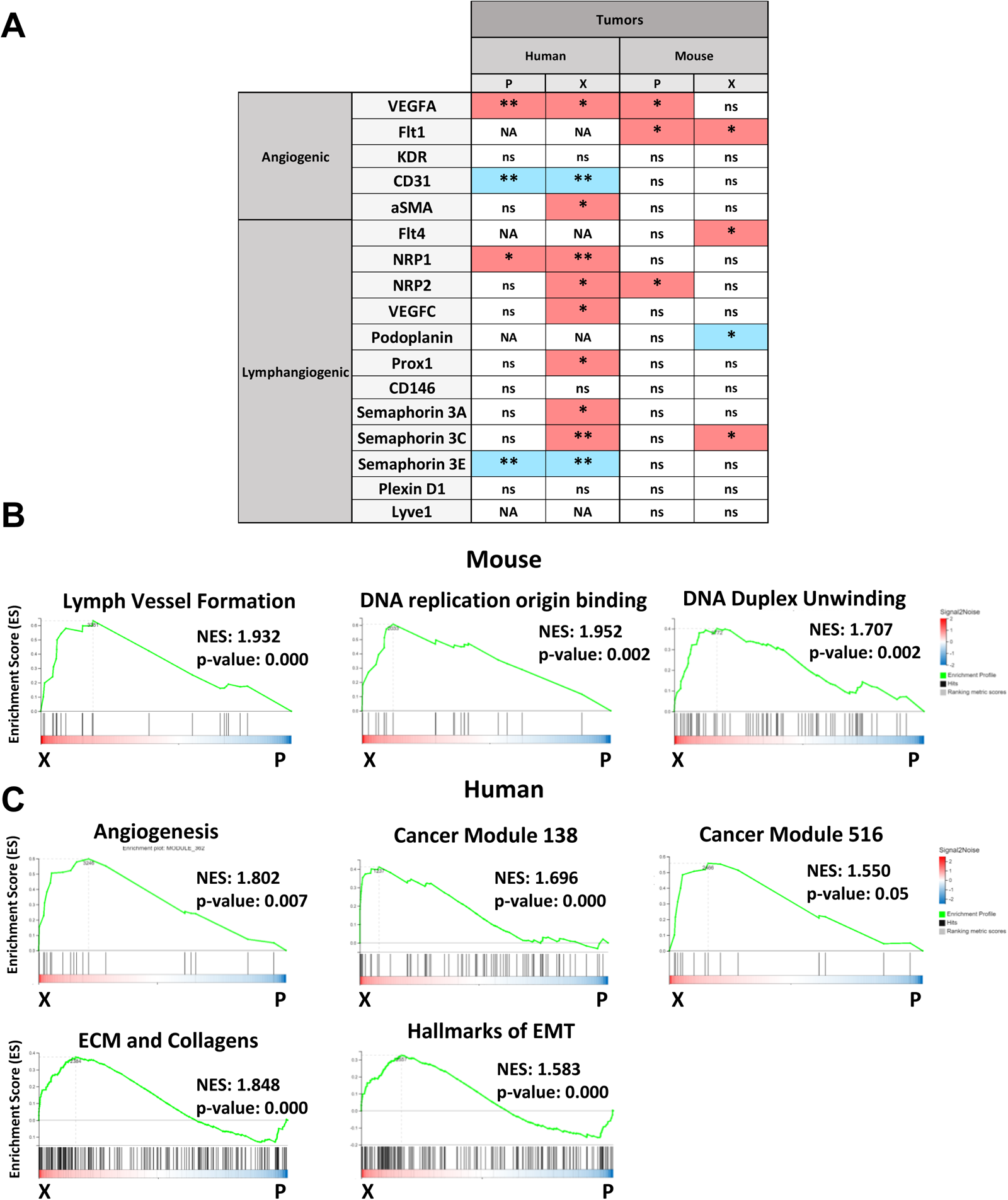
X multi-irradiated tumors have a higher angiogenic, lymphangiogenic, and EMT transcriptome profile. **(A)** Expression of angiogenic and lymphangiogenic genes in the human and mouse transcriptomes of MDAMB231 P and X multi-irradiated tumors compared to control tumors. Red indicates upregulation of the gene compared to the respective control, while blue represents downregulation. NA (non-applicable). GSEA Enrichment Analysis of Pathway Clusters in mouse **(B)** and human **(C)** transcriptome. This GSEA enrichment graphs illustrate the pathway clusters that are significantly over-represented in the X tumors compared to P tumors. GSEA was performed using the GSEA software from the Broad Institute on a ranked list of genes based on differential expression between X and P tumors. Pathways were grouped into clusters using hierarchical clustering based on their functional similarity. The results are presented as the mean of at least three independent experiments ± SD. Statistical analysis was performed using one-way ANOVA to compare differences between the control and irradiated groups (P or X). Statistical significance is denoted as follows: *P < 0.05, **P < 0.01, ***P < 0.001, ****P < 0.0001, NS (non-significant).

Examining the effect of P and X irradiation on tumors formed *in vivo* at the protein level revealed distinctly different protein profiles among control, P, and X tumors. This diversity in protein profiles enhances our understanding of the unique molecular characteristics and behaviors of P tumors compared to X tumors (**Figure 9A**). Analyzing the highest and lowest deregulated proteins in P and X tumors showed minimal overlap between the two groups (**Figure 9B**); SNCG (gamma-synuclein protein) which plays a multifaceted role in BC by promoting tumor progression, enhancing cell proliferation, and facilitating metastasis [27], CPD (Carboxypeptidase D protein), whose expression is increased in BC and can stimulates the production of nitric oxide, promoting the survival of breast and prostate cancer cells [28, 29]; and ARG2 (Arginase 2 protein), which is important in T-cell depletion and breast cancer brain metastasis [30], are commonly downregulated between P and X irradiations. Conversely, PLG (plasminogen), a key player in modulating the tumor microenvironment and progression of BC [31]; BCS1L (C1 (ubiquinol-cytochrome c reductase) synthesis-like (BCS1L)) which is implicated in cancer-related fatigue [32]; and TOP2B (topoisomerase 2B), a marker of proliferation in BC [33] and is down-regulated by trastuzumab, the reference treatment for HER2-positive BC [34], are commonly upregulated in both P and X groups. The complete list of up and downregulated proteins is in **Table S2.** Consistent with this finding, the Principal Component Analysis (PCA) plot of these tumors also depicted distinct clusters based on protein expression profiles. The separate clustering of P and X tumors supports the notion that they exhibit different molecular features and behaviors. Furthermore, the PCA plot indicated that the cluster of P tumors is closer to the control tumors than the cluster of X tumors, suggesting a higher similarity in protein expression patterns between P tumors and control tumors compared to X tumors (**Figure 9C**).

**Figure 9.**
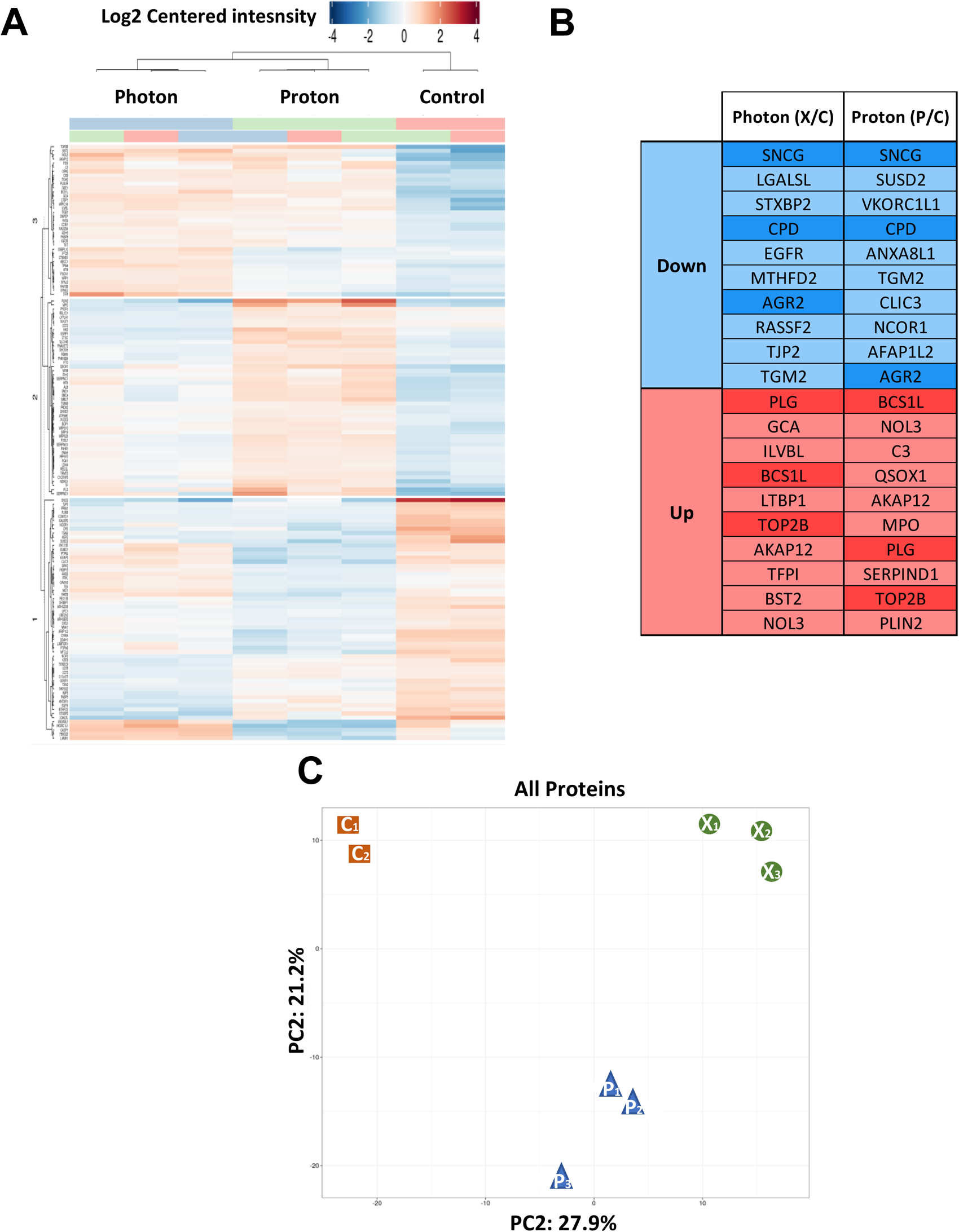
P and X multi-irradiated tumors have distinct proteome profiles. **(A)** The heatmap representation of all significant proteins (rows) in all P and X multi-irradiated tumors compared to control tumors (columns). **(B)** Ten lowest (blue) and highest (red) deregulated proteins in proteome analysis of X and P multi-irradiated tumors compared to control tumors. Dark blue and dark red indicated the common deregulated proteins. **(C)** PCA (Principal Component Analysis) of protein expression in P and X Tumors. The PCA plot visualize the differences in protein expression profiles between P and X tumors. Each point represents a tumor sample: Red dots represent control tumors, Blue dots represent P tumors, Green dots represent X tumors. This PCA plot shows a clear separation between P and X tumors.

### Anti-VEGFC antibodies exhibit significant therapeutic efficacy in TNBC tumor models

*In vitro* experiments underscored the crucial role of VEGFC in driving the aggressiveness of TNBC. Consequently, we investigated the therapeutic impact of targeting VEGFC in these experimental models. Mice injected with highly aggressive MDAMB231 cells were treated twice a week with either isotype control or anti-VEGFC antibodies (7.5 mg/kg).

Remarkably, even at this relatively low dose compared to the dose used for the treatment of experimental prostate tumors with these antibodies (40 mg/kg, as described in patent WO2011127519A1), anti-VEGFC antibodies profoundly delayed the growth of experimental TNBC tumors, with some tumors exhibiting nearly complete regression (**Figure 10A**). Moreover, tumor volume and weight were significantly reduced at the end of the experiment (**Figure 10B-D**). Considering the substantial increase in VEGFC expression post-irradiation with both P and X, combining irradiation with P or X could potentially enhance the therapeutic effects of anti-VEGFC treatment.

**Figure 10.**
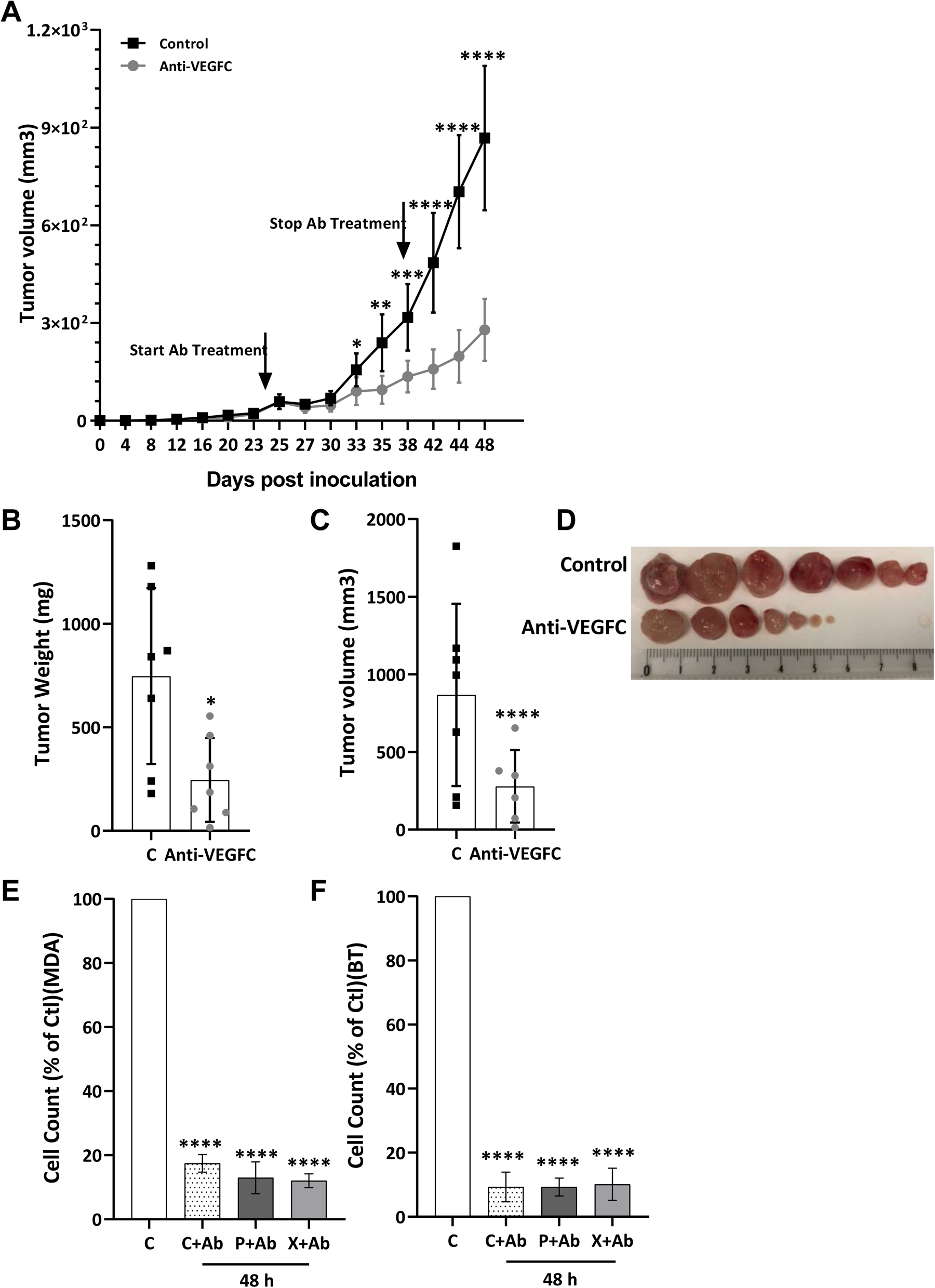
Anti-VEGFC antibodies slow down the growth of experimental TNBC. The evaluation of tumors generated following xenografting of MDAMB231 cells in immunodeficient mice. Starting from day 25^th^ post injection of 5*10^6^ tumor cells subcutaneously, half of the mice received anti-VEGFC antibody (7.5 mg/kg twice a week), while the other half received an irrelevant antibody (Control), for a total duration of 18 days. **(A)** Monitoring of tumor growth in the two groups over a period of 48 days. **(B)** Measurement of tumor weights on the last day of the experiment for both groups. **(C)** Measurement of tumor volumes on the last day of the experiment for both groups. **(D)** Representative image of the tumor xenografts showcasing the morphology and growth of the tumors in each experimental group. **(E-F)** The effect of anti-VEGFC antibody on MDAMB231 and BT549 multi-irradiated for 48h. Differences between the control and treated groups were analyzed using the Student’s t-test **(A-C**) and one-way ANOVA **(E-F)**. Statistical significance was denoted as follows: *P < 0.05, **P < 0.01, ***P < 0.001, ****P < 0.0001, NS (non-significant).

To further investigate this, the effect of anti-VEGFC antibody was studied on P and X multi-irradiated MDAMB231 and BT549 cells. Similar effects were observed across all conditions: anti-VEGFC antibody decreased proliferation by up to 90% in both control and multi-irradiated MDAMB231 and BT549 cells (**Figure 10E-F**).

These findings underscore the critical role of VEGFC in promoting proliferation, particularly in the context of irradiated TNBC cells. Targeting VEGFC with an antibody could potentially mitigate the enhanced proliferative capacity induced by both P and X irradiation. This approach suggests a promising therapeutic strategy to suppress TNBC progression following radiotherapy whether with P or X, by counteracting their potential detrimental effects.

## Discussion

TNBC represents a significant unmet medical need, with low survival rates post-diagnosis. Improving outcomes for this devastating disease remains a paramount challenge. While advances in radiotherapy have transformed patient management, individual responses to treatment vary widely. Our research has spotlighted VEGFC as a critical driver of TNBC aggressiveness, with implications that extend to other tumor types in response to radiotherapy. The conventional belief surrounding P-based radiotherapy centers on its precision, aimed at sparing healthy tissue. However, our provocative theory posits that radiotherapy exerts both anti-and pro-tumor effects, notably by modulating the tumor microenvironment. Our experiments partially confirm this hypothesis, revealing that radiotherapy stimulates the expression of key factors mediating lymphatic vessel development, thereby enhancing the potential for tumor cell dissemination via the lymphatic network.

However, this assumption must be tempered, as lymphatic vessels also facilitate the recruitment of immune cells to the tumor site, promoting anti-tumor immune responses. The delicate balance between these opposing effects underscores the need for caution when considering therapies that may induce VEGFC expression and lymphatic vessel development [35].

The crucial question lies in identifying the therapeutic window that balances tumor cell destruction with the inhibition of VEGFC-mediated lymphatic vessel development. Additionally, we must determine the optimal timing and administration of radiotherapy, chemotherapy, and targeted therapies. While textbooks depict P as more efficient than X, the simplistic calculation of the Effective Biologic Ratio (EBR) may not accurately reflect clinical outcomes, particularly considering patients receive multiple rounds of irradiation.

How and when radiotherapy could be the most efficient, and how and when chemotherapeutic agents and/or targeted therapies can be administered? Which kind of radiotherapy is the most appropriate and does VEGFC targeting can be reliable? In the textbooks, P appears as most efficient than X considering the EBR results of 1.1 in favor of P. But how this EBR is calculated? Generally, it is calculated following only one irradiation of tumor cells at increasing doses and then count the number of resistant clones. By this way, 1.1 less clones are observed with P than X. This simple method is too “naïve” to be translated to patients as they received multiple rounds of irradiation to eradicate remaining tumor cells. Therefore, different possibility of adaptation and shaping of the normal tissue can lead to a favorable “milieu” that will feed the possible remaining tumor cells that are also capable of genetic plasticity and adaptation.

Our findings suggest that P and X radiotherapy induce distinct modifications in TNBC tumor cells and the tumor microenvironment, resulting in different tumor characteristics *in vivo*. This study reveals, for the first time, that while P multi-irradiated TNBC cells (modeled to simulate resistant cells to radiotherapy) lead to larger tumors compared to X multi-irradiated cells, the protein and transcriptome profiles indicate higher aggressiveness in X multi-irradiated tumors. These observations imply that the long-term effects of X-based radiation could potentially contribute to more aggressive relapses, particularly if resistant cells persist. This highlights the importance of understanding the specific molecular and phenotypic changes induced by different radiation types to optimize treatment strategies and improve outcomes for TNBC patients. Future research should focus on elucidating the mechanisms underlying these differences and developing targeted interventions to mitigate the risks associated with X-based radiotherapy in TNBC. Comparative clinical trials are underway to provide further insights into the effects of X versus P radiation.

Our findings underscore the importance of considering tumor-specific factors, such as basal VEGFC expression levels and the potential induction of VEGFC post-irradiation, in treatment planning. Our research highlights the promising therapeutic potential of anti-VEGFC antibodies, which are readily available for clinical trials.

Moving forward, our focus will be on defining the optimal therapeutic schedule, including the possibility of administering anti-VEGFC antibodies before irradiation to mitigate radiation-induced stress responses. Based on the favorable outcomes observed in our aggressive TNBC model, we eagerly anticipate the opportunity to evaluate such treatments in future clinical trials.

## Acknowledgments

This work was performed using the microscopy (PICMI), cytometry (CYTOMED) and mouse facilities of IRCAN. The materials of CytoMed were supported by the Conseil Général 06, the FEDER, the Ministère de l’Enseignement Supérieur, the Région Provence Alpes-Côte d’Azur and the INSERM.

## Supplement Figure Legends

**Figure S1.**
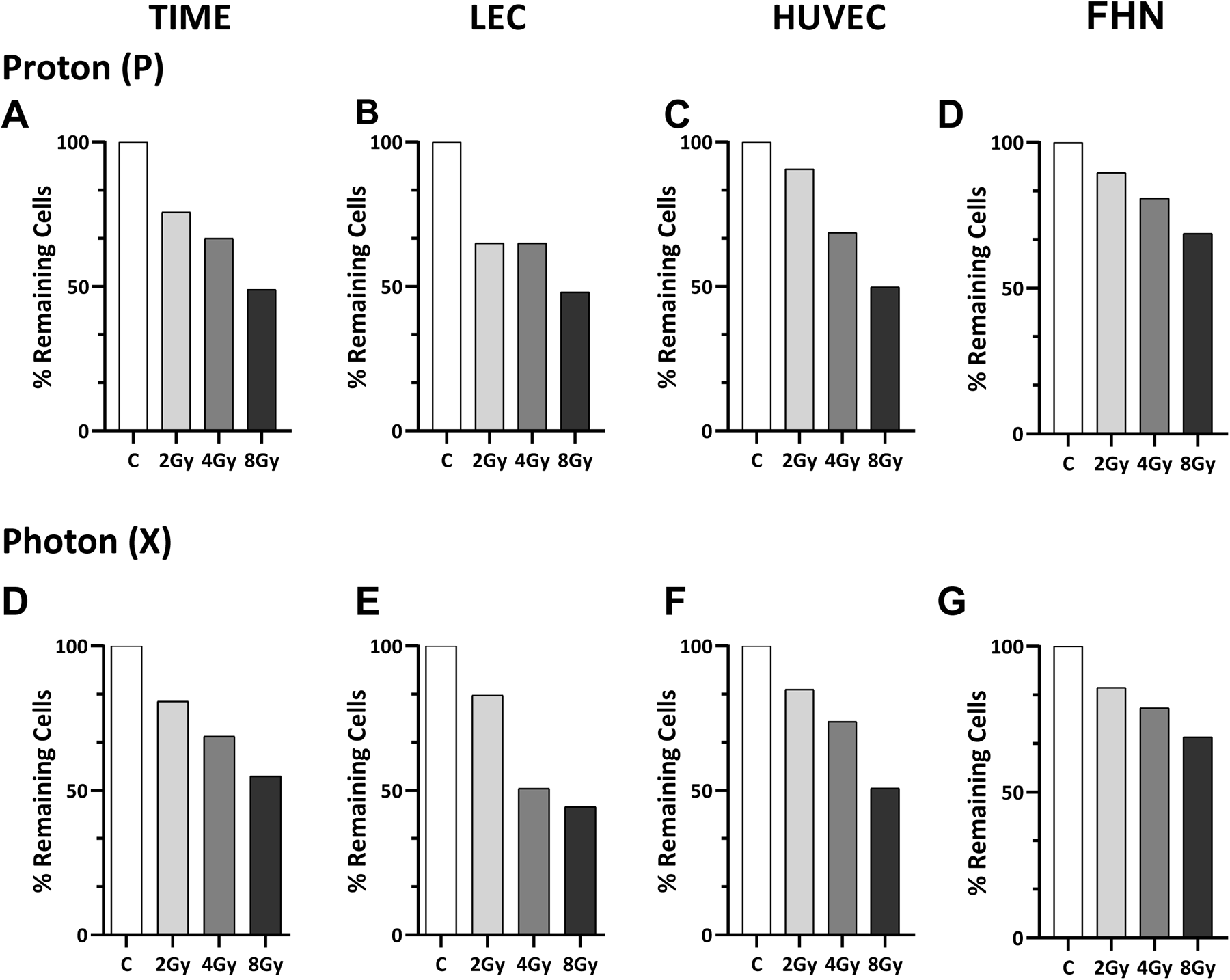
Remaining cells 48 h post P and X irradiations in TIME, LEC, HUVEC and FHN cells. Remaining cells post 2, 4 and 8 Gy Proton **(A-D)** and photon **(E-G)** irradiations compared to corresponding controls in Time, LEC and HUVEC cells.

**Figure S2.**
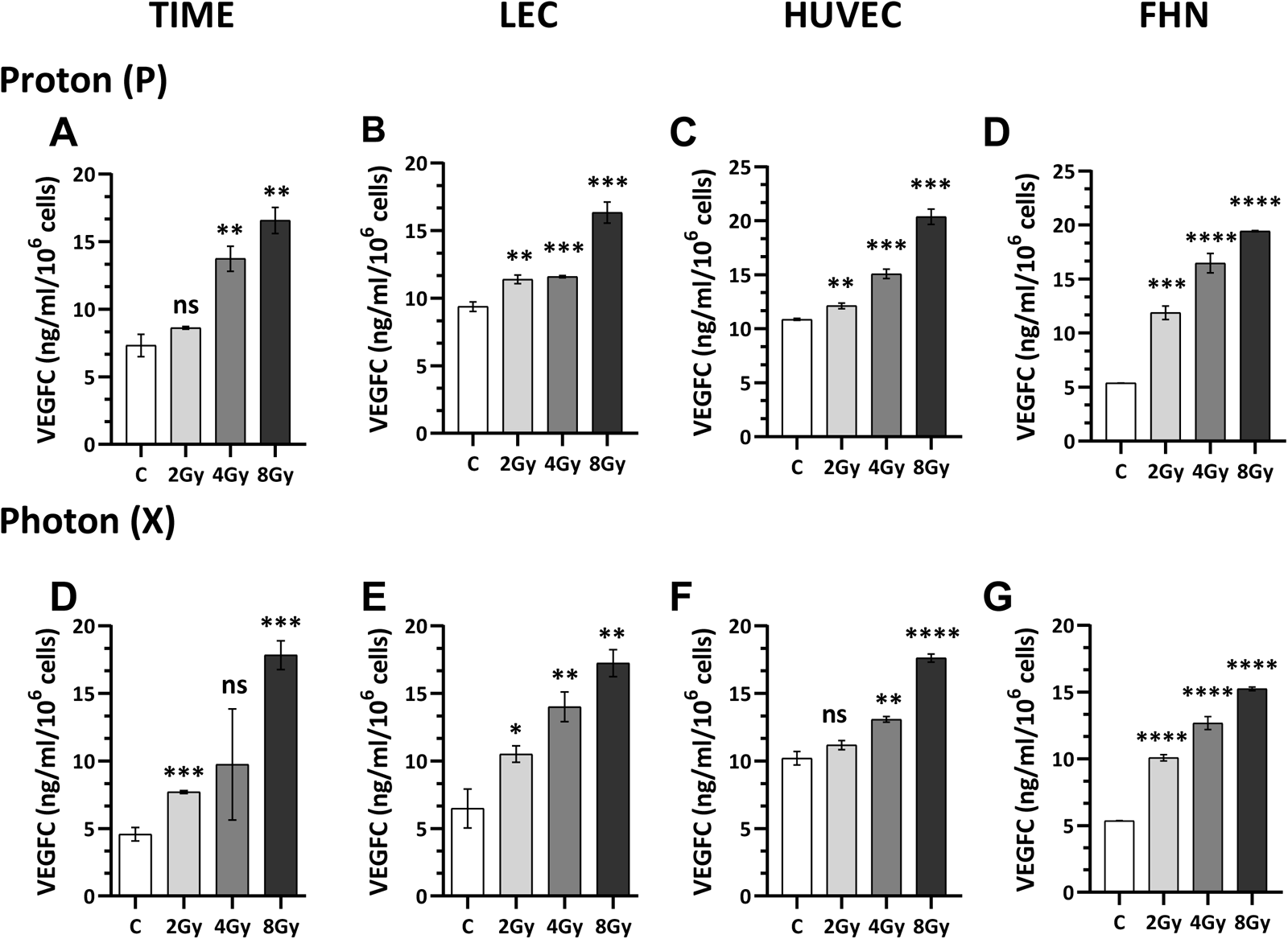
VEGF expression 48 h post P and X irradiations in TIME, LEC, HUVEC and FHN cells. Quantification of secreted VEGFC protein levels in supernatant of single 2, 4 and 8 Gy P **(A-D)** and X-irradiated **(E-G)** TIME, LEC and HUVEC cells 48 hours post irradiation compared to respective controls (C) using ELISA.

**Figure S3.**
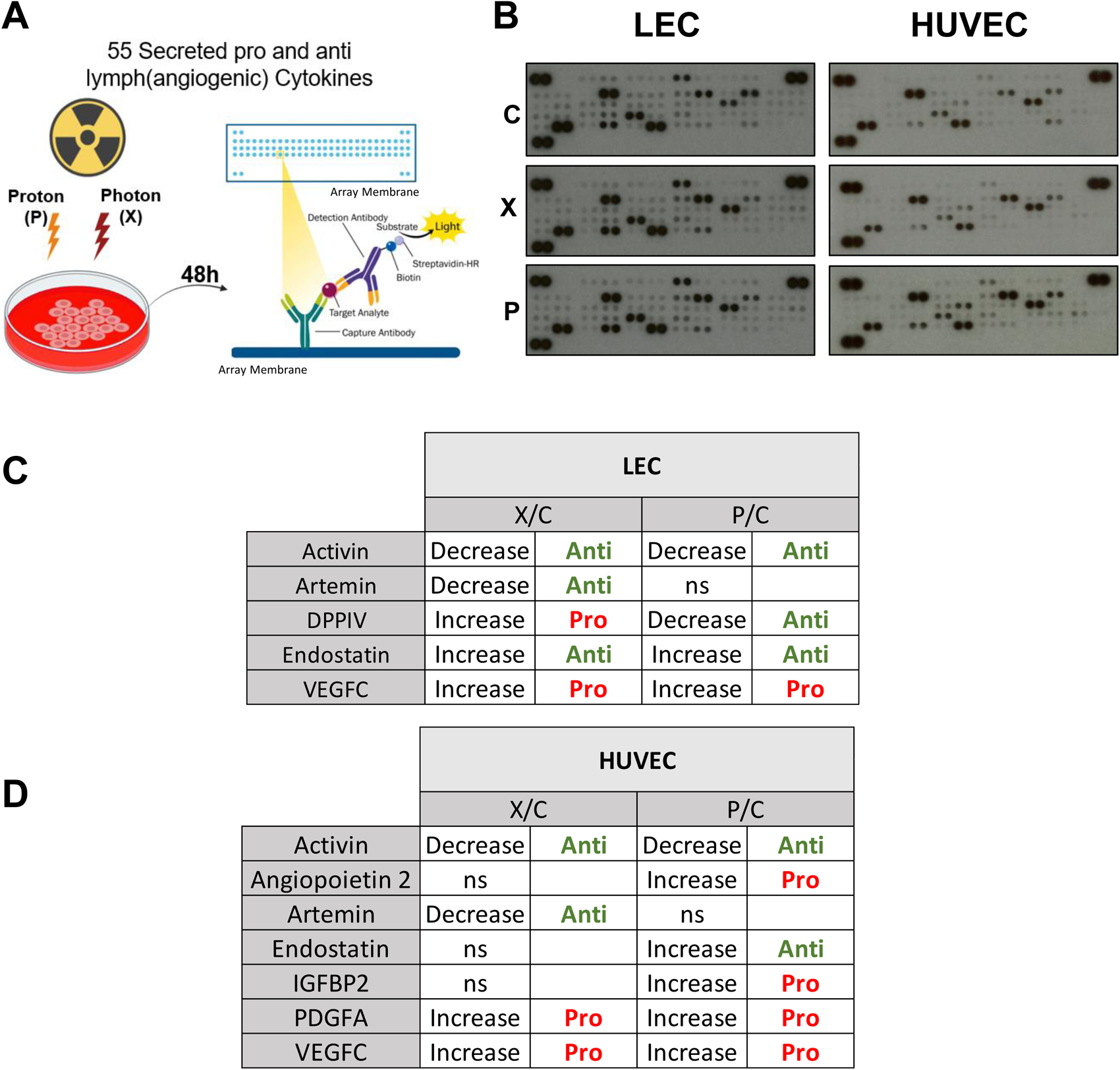
(Lymph)Angiogenic proteome profile of LEC and HUVEC cells. **(A-B)** Analysing the expression profiles of (lymph)angiogenesis-related proteins in LEC and HUVEC supernatants 48 hours post P and X irradiations. **(C-D)** Significant alterations in the secretion of (lymph)angiogenic proteins were observed in the supernatants of LEC and HUVEC 48 hours after exposure to P and X irradiations. The secretion levels of each cytokine were compared to their respective controls and expressed as either increased or decreased compared to the control. ‘Anti’ refers to Anti-(lymph)angiogenic proteins, while ‘Pro’ indicates Pro-(lymph)angiogenic.

**Figure S4.**
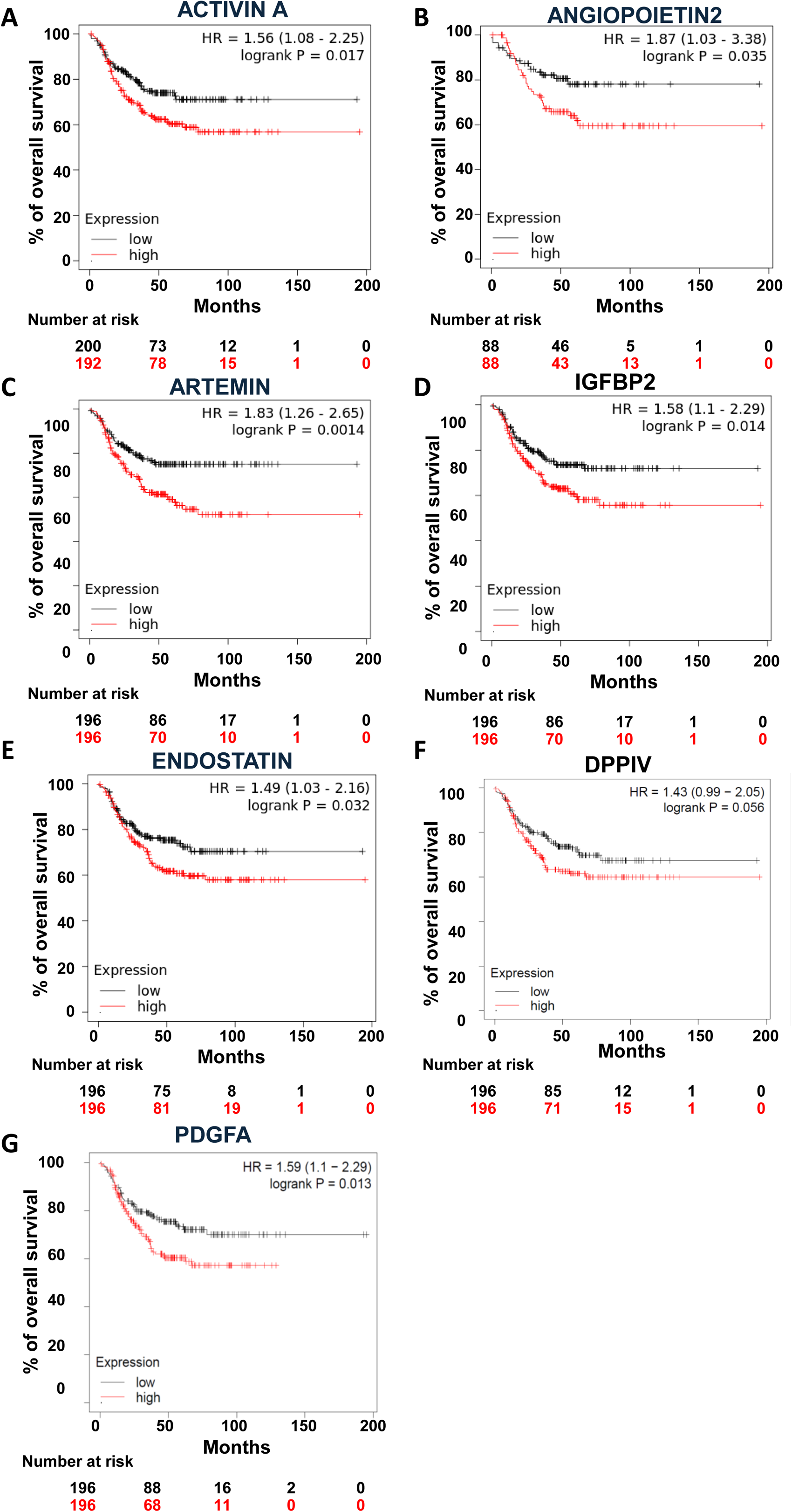
Survival profile of TNBC patients. Analysis of overall survival (OS) of TNBC patients using the Kaplan Meier softaware (https://kmplot.com/analysis/). OS was calculated from patient subgroups with mRNA levels of **(A)** Activin A **(B)** Angiopoietin2 **(C)** Artemin **(D)** IGFBP2 **(E)** Endostatin **(F)** DPPIV **(G)** PDGFA that were less or greater than the best cut-off value.

**Figure S5.**
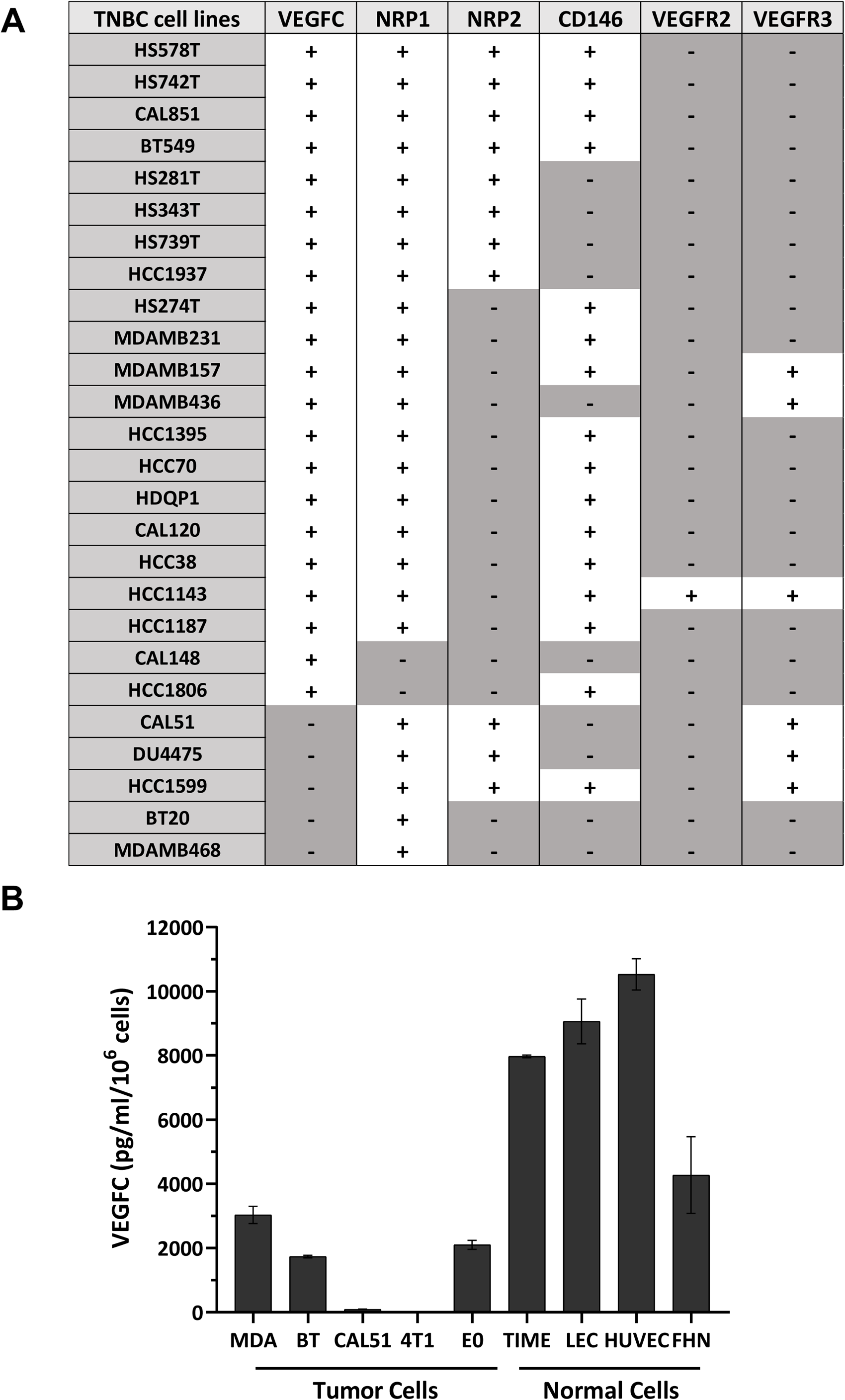
Relative expression of VEGFC in TNBC cells and normal cells. **(A)** VEGFC pathway component expression in TNBC cell lines. **(B)** Quantification of secreted VEGFC protein levels in supernatant of MDAMB231, BT549, CAL51, 4T1, E0, TIME, LEC, HUVEC and FHN cells using ELISA.

**Figure S6.**
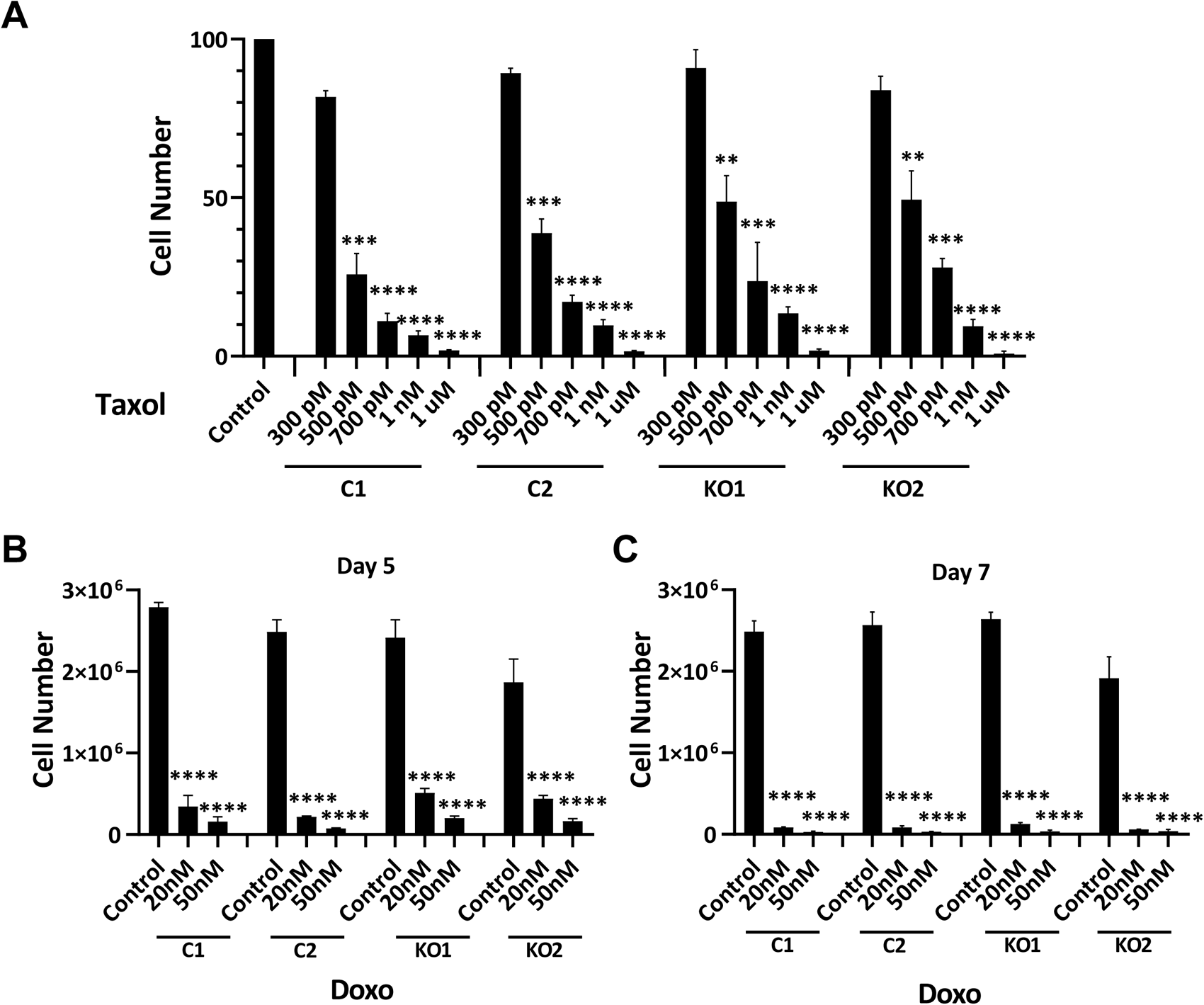
The effect of chemotherapy on VEGFC KN MDAMB231. (**A**) Assessment of remaining CRISPR/Cas9-edited MDAMB231 cells, with the empty vector serving as the control (C) and VEGFC knockout (KO) as the experimental condition, 48 hours following Taxol treatment during a week in different concentrations compared to their respective nontreated controls. **(B-C)** The clonogenic potential of the same cells treated with Taxol was evaluated. **(D-E)** Cell count was performed on the same cells This assessment was performed five days post-treatment with Doxorubicin at various dosages. Two clones from the control group and two clones from the KO group were evaluated for each condition. The results are presented as the mean of at least three independent experiments ± SD. Statistical analysis was performed using one-way ANOVA to compare differences between the control and irradiated groups (P or X). Statistical significance is denoted as follows: *P < 0.05, **P < 0.01, ***P < 0.001, ****P < 0.0001, NS (non-significant).

**Table S1.**
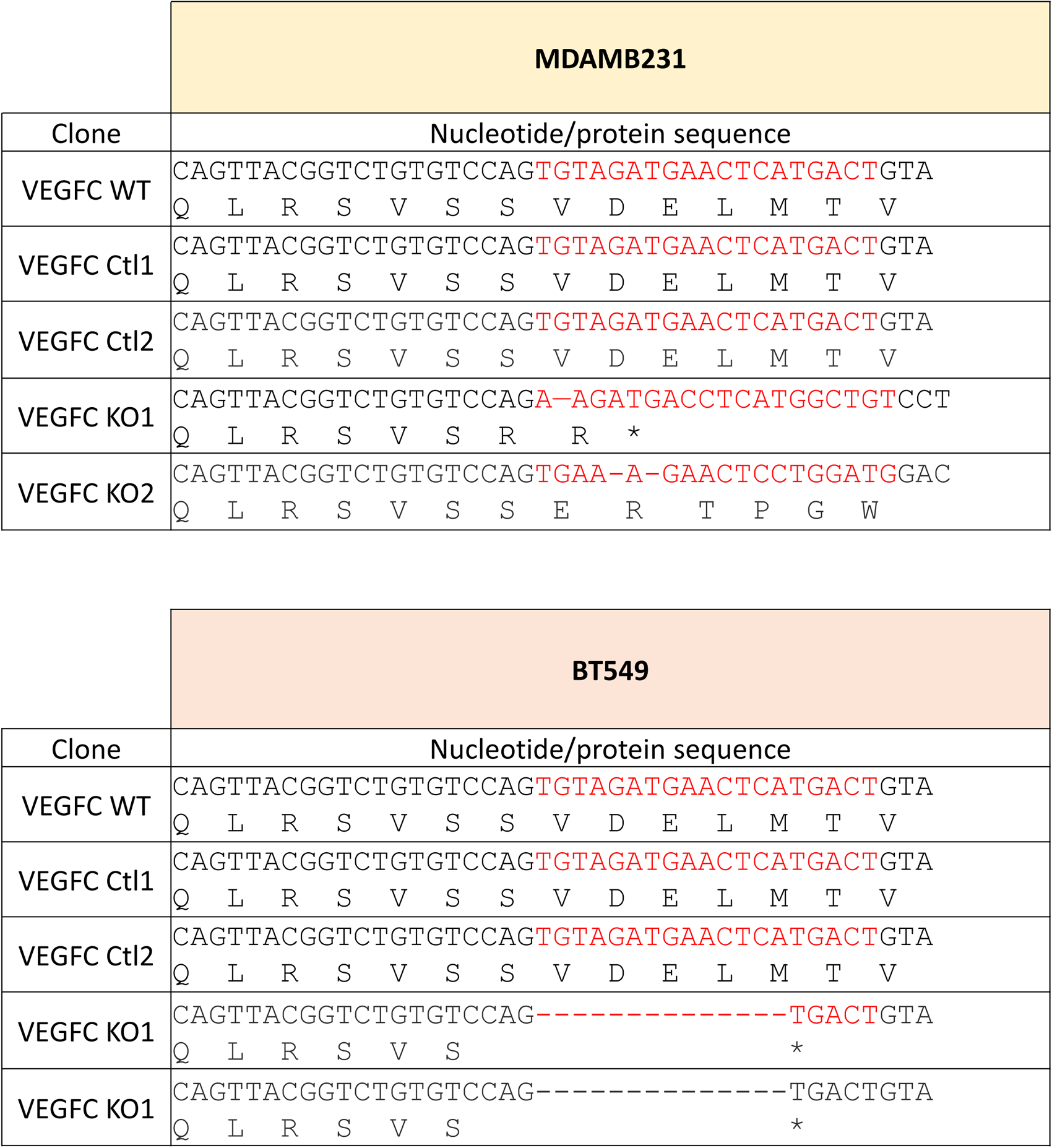
VEGFC invalidation by CRISPR-Cas9 for MDAMB231 and BT549. Wild-type, fake-mutated (Ctl) and mutated sequences. *: Stop codon.

**Table S2.**
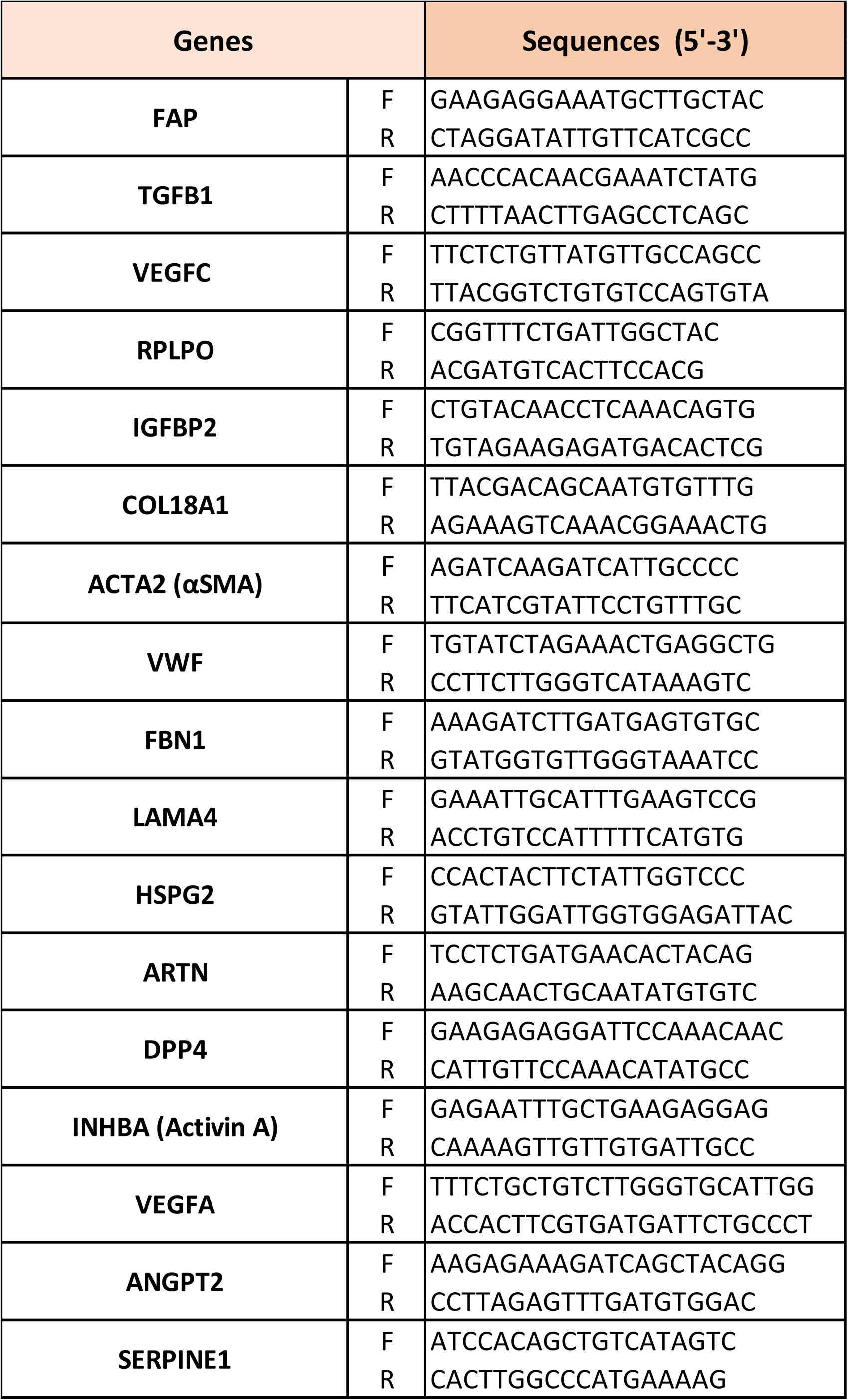
Primer Sequences.

**Table S3.**
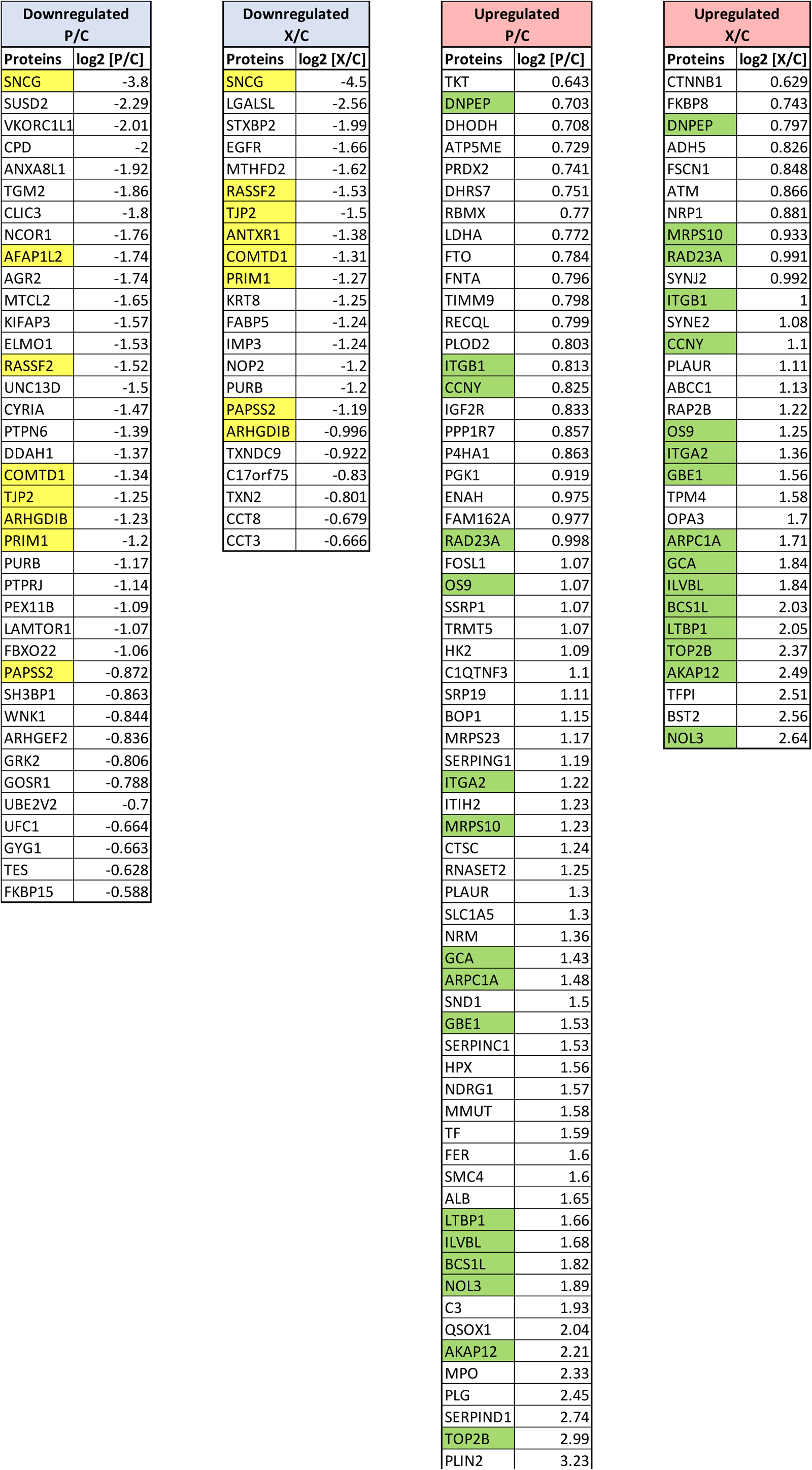
List of up and downregulated proteins based on proteome analysis of the mice tumors. The common downregulated proteins in both P and X tumors are highlighted with yellow. The common upregulated proteins between P and X tumors highlighted with green.

